# Single-molecule imaging reveals control of parental histone recycling by free histones during DNA replication

**DOI:** 10.1101/789578

**Authors:** Dominika T. Gruszka, Sherry Xie, Hiroshi Kimura, Hasan Yardimci

## Abstract

Faithful replication of chromatin domains during cell division is fundamental to eukaryotic development. During replication, nucleosomes are disrupted ahead of the replication fork, followed by their rapid reassembly on daughter strands from the pool of recycled parental and newly synthesized histones. Here, we use single-molecule imaging and replication assays in *Xenopus laevis* egg extracts to determine the outcome of replication fork encounters with nucleosomes. Contrary to current models, the majority of parental histones are evicted from the DNA, with histone recycling, nucleosome sliding and replication fork stalling also occurring but at lower frequencies. The anticipated local histone transfer only becomes dominant upon depletion of free histones from extracts. Our studies provide the first direct evidence that parental histones remain in close proximity to their original locus during recycling and reveal that provision of excess histones results in impaired histone recycling, which has the potential to affect epigenetic memory.

## INTRODUCTION

Eukaryotic genomes are organized into chromatin, which influences many cellular processes, ranging from DNA replication and repair to gene transcription. The basic unit of chromatin is a nucleosome, which consists of 145–147 base pairs of DNA wrapped around an octameric histone protein core, formed from two copies of each of the four histones: H2A, H2B, H3 and H4. Histones H3 and H4 assemble into a symmetric hetero-tetramer and the two H2A–H2B dimers are docked onto the (H3– H4)_2_ tetramer (Luger et al., 1997). Nucleosomes are very stable nucleoprotein complexes but they are also highly dynamic with regards to their conformation, composition and positioning within chromatin (Lai and Pugh, 2017; Zhou et al., 2019). Nucleosome dynamics control DNA accessibility and are regulated by complex interplay of numerous factors, such as chromatin remodelers, histone chaperones, modifying enzymes and polymerases (Lai and Pugh, 2017).

Chromatin is partitioned into domains, which either promote or block transcription, and hence determine the cellular identity. Nucleosomes in transcriptionally active and silenced chromatin domains carry specific histone post-translational modifications (PTMs) and/or distinct histone sequence variants. The disordered tails of histones H3 and H4 are primary targets for PTMs associated with different chromatin states; for example, tri-methylation of histone H3 at lysine 36 (H3-K36Me3) and acetylation of histone H4 at lysine 16 (H4-K16Ac) mark transcriptionally active chromatin, whereas tri-methylation of histone H3 at lysine 9 and 27 (H3-K9Me3 and H3-K27Me3) tag transcriptionally silenced chromatin domains (Reinberg and Vales, 2018; Stillman, 2018). Therefore, maintenance of cellular identity through mitotic cell division relies on faithful transfer of information encoded in both DNA sequence (genetic inheritance) and nucleosome landscape (epigenetic inheritance). Semiconservative DNA replication ensures genetic inheritance, but it presents a major challenge to chromatin, which undergoes significant structural reorganization, starting from disassembly of parental nucleosomes and ending in restoration of nucleosome landscape on daughter strands (Alabert et al., 2017; MacAlpine and Almouzni, 2013; Ramachandran and Henikoff, 2015; Serra-Cardona and Zhang, 2018). The molecular mechanisms of replication-coupled epigenetic inheritance are poorly understood.

In order to allow parental DNA unwinding and subsequent nascent strand synthesis, each and every nucleosome must be transiently disrupted ahead of the replication fork. Nucleosome destabilization is localized to an average of two nucleosomes immediately ahead of the replication fork (Gasser et al., 1996) and leads to release of (H3–H4)_2_ tetramer and H2A–H2B dimers from the DNA (Jackson, 1990; Xu et al., 2010). It remains unclear what molecular forces trigger the localized nucleosome eviction but a number of physical and chromatin factors have been implicated in this process, including unzipping (Shundrovsky et al., 2006) and positive supercoiling (Gupta et al., 2009) of the DNA, physical collision between the nucleosome and the replisome (Sogo et al., 1986), and histone chaperone complex FACT (Foltman et al., 2013; Kurat et al., 2017).

After passage of the replication fork, nucleosomes are rapidly reassembled on the two daughter strands, from the pool of recycled parental and newly synthesized histones. Nucleosome reassembly starts with the deposition of (H3–H4)_2_ tetrameric histone core, followed by the association of two H2A–H2B dimers (Alabert et al., 2017; MacAlpine and Almouzni, 2013; Ramachandran and Henikoff, 2015). Current models suggest that nucleosomes deposited on newly replicated DNA contain either parental or new H3–H4 histones (with the exception of nucleosomes containing H3.3 variant). This implies two distinct replication-coupled nucleosome assembly pathways on nascent DNA: the transfer (recycling) of parental histones released from nucleosomes disrupted by the replisome passage and *de novo* deposition of newly synthesized histones. Nucleosomal H2A–H2B dimers are more dynamic than (H3–H4)_2_ tetramers and readily exchange with the pool of newly synthesized histones throughout the cell cycle (Annunziato, 2013; Jackson, 1990; Kimura and Cook, 2001; Louters and Chalkley, 1985; Thiriet and Hayes, 2006). Consequently, old and newly synthesized H2A–H2B dimers can form nucleosomes with both parental and new (H3–H4)_2_ tetramers (Annunziato, 2015; Xu et al., 2010).

Quantitative proteomics studies indicate that nucleosomes deposited on newly replicated DNA are composed of approximately equal amounts of new and old H2A, H2B, H3 and H4 histones (Alabert et al., 2015), implying that all parental histones are fully recycled during replication. It has also been reported that parental histones are recycled with PTMs (Alabert et al., 2015) and that their genomic localization, whether in active or repressed chromatin, is preserved on daughter strands through histone recycling (Madamba et al., 2017; Reveron-Gomez et al., 2018). However, recent studies using mouse embryonic stem cells demonstrated that while repressed chromatin domains are indeed preserved through the local re-deposition of parental H3–H4 histones at the replication fork, parental H3–H4 histones associated with active chromatin domains did not exhibit such preservation (Escobar et al., 2019). Furthermore, fluorescence imaging-based analysis of parental H3 histone recycling in HeLa cells over two cell divisions revealed rates of parental histone loss that were higher than the expected 50% per cell cycle (Clement et al., 2018).

Recent studies into the mechanism of parental histone segregation onto replicating DNA showed that histones H3–H4 distribute more or less equally between the two strands; depending on the study, a weak bias was observed towards either the leading (Petryk et al., 2018) or lagging strand (Yu et al., 2018). Importantly, several replisome components are involved in parental histone segregation. The N-terminal domain of MCM2, a component of the CMG replicative helicase, contains a histone H3–H4-binding domain (Huang et al., 2015; Ishimi et al., 1998) and promotes the transfer of parental H3–H4 to the lagging strand (Petryk et al., 2018), in association with the Ctf4 adaptor protein and Pol *α* (Gan et al., 2018). Dpb3 and Dpb4, two non-essential subunits of yeast Pol *ε*, facilitate the parental H3–H4 transfer to the leading strand (Yu et al., 2018). In addition, various histone chaperones, chromatin factors and other replisome components have been implicated in parental histone recycling and/or the inheritance of chromatin states (Alabert et al., 2017; Serra-Cardona and Zhang, 2018). Taken together, these findings support a mechanism for replication-coupled parental histone recycling whereby, upon eviction from the DNA, parental histones H3–H4 are retained close to the replisome through a series of protein-protein interactions, resulting in their targeted and localized re-deposition behind the replication fork.

To date, most studies looking into the mechanisms of replication-coupled parental histone recycling employ bulk and/or steady-state approaches, which lack the spatial and temporal resolution needed to unravel the molecular detail of this highly dynamic process. Here, we report a real-time single-molecule imaging platform that utilizes microfluidics-based DNA replication in *Xenopus laevis* egg extracts, protein engineering and TIRF microscopy to gain mechanistic insight into the outcome of replication fork collision with nucleosomes. Our approach allows simultaneous visualization of parental histones and replication forks as they navigate through the nucleosomal environment of individual DNA molecules.

## RESULTS

### Assembly of fluorescent nucleosomes on λ DNA

Building on the single-molecule methodology developed by (Loveland et al., 2012), which allows real-time imaging of growing replication bubbles in *Xenopus* egg extracts, we wanted to visualize fork collisions with nucleosomes. To be able to track nucleosomes during replication, we labelled *Xenopus laevis* histones H2A, H2B, H3 and H4 with small fluorescent dyes using thiol-modifications of engineered cysteines (Figures 1A and 1B). Histones were expressed and purified from *Escherichia coli* that did not carry any post-translational modifications, as verified by mass spectrometry (Table S1). To ensure that our observations are representative of nucleosome dynamics, we labelled all four histones, at various positions in their structure, using different fluorophores. Histones H2A and H2B were labelled at their C-termini; H2A was labelled with Cy5 at position 119 (H2A-K119C^Cy5^; 66% labelling efficiency) and H2B was labelled with AlexaFluor647 at position 112 (H2B-T112C^A647^; 45% labelling efficiency). Histone H3 was labelled with Cy5 at position 36 (H3-K36C^Cy5^; near 100% labelling efficiency), in the disordered tail, and with AlexaFluor647 at position 80 (H3-T80C^A647^; 30% labelling efficiency), in the folded histone core. Histone H4 was labelled with AlexaFluor647 at position 63 (H4-E63C^A647^; 95% labelling efficiency), within the histone fold (Figures 1A and 1B).

**Figure 1.**
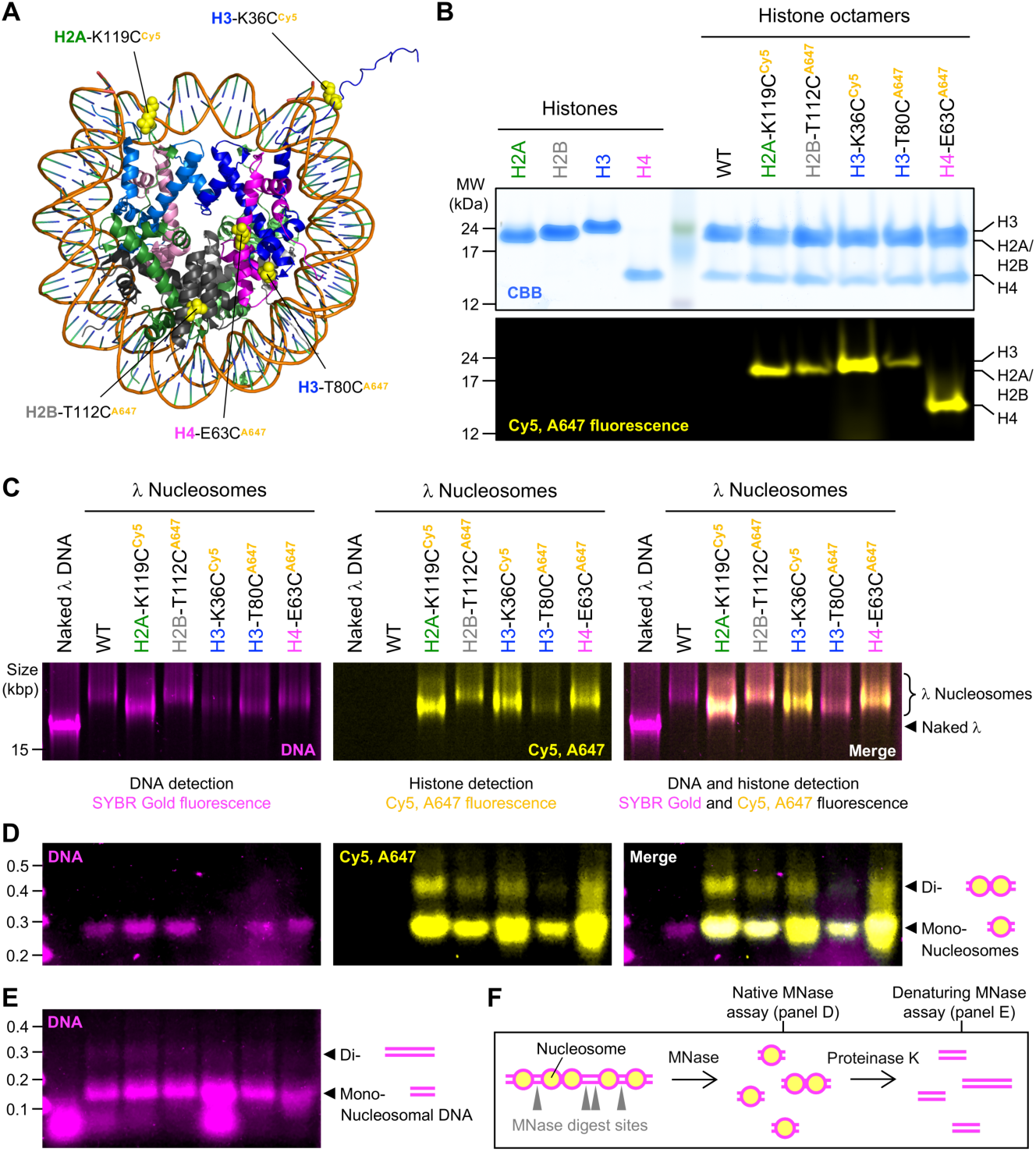
Assembly of fluorescent nucleosomes on λ DNA. (A) Crystal structure of the *Xenopus* nucleosome (PDB 1AOI) illustrating the location and type of fluorescent dye (Cy5 or AlexaFluor647 – abbreviated as A647) used to label histones. Histones are color-coded (H2A – green, H2B – grey, H3 – blue and H4 – magenta) and the two chains of the same histone type can be distinguished by different color shading. For clarity, only one of the two histones of the same type is marked and labelled. (B) SDS-PAGE analysis of wild-type (WT) and fluorescently-labelled histones and histone octamers. Top panel shows Coomassie Brilliant Blue (CBB) staining whereas bottom panel illustrates fluorescence signal of histones labelled with Cy5 or AlexaFluor647. (C) Electrophoretic mobility shift assay (EMSA) for WT and fluorescently-labelled nucleosomes reconstituted on *λ* DNA. Left panel shows SYBR Gold staining of the DNA (magenta), central panel shows Cy5 and AlexaFluor647 fluorescence signal (yellow) of labelled histones and right panel is the composite of both detection modes. Naked *λ* DNA (∼48.5 kbp, first lane) migrates through 0.5 % agarose faster than nucleosomal *λ* templates, containing either WT or fluorescently-labelled histones. (D) Native micrococcal nuclease (MNase) protection assay for WT and fluorescently-labelled nucleosomes reconstituted on *λ* DNA. MNase preferentially digests unprotected DNA in linker regions between nucleosomes (see also panel F). Products of MNase digest were resolved in 1.5 % agarose under native conditions revealing intact mono- and di-nucleosomes for nucleosomal templates and complete digest of naked *λ* DNA (first lane). Signal detection as in panel C. (E) Denaturing micrococcal nuclease (MNase) protection assay for WT and fluorescently-labelled nucleosomes reconstituted on *λ* DNA. Here, products of MNase digest were first deproteinated with proteinase K (see also panel F) in the presence of SDS and then resolved in 1.5 % agarose, yielding DNA fragments protected by mono-(∼150 bp band) and di-nucleosomes (∼300 bp band) for nucleosomal templates, and short (<100 bp) fragments for naked *λ* DNA (first lane). (F) Schematic overview of native and denaturing MNase protection assays.

Histone octamers, containing one of the four histones labelled fluorescently, were then used to reconstitute nucleosomes on biotinylated *λ* DNA (Figure 1). Nucleosome reconstitution was carried out by NaCl gradient dialysis, which recapitulates thermodynamically favorable binding distributions (Sekinger et al., 2005; Thastrom et al., 2004a; Thastrom et al., 2004b; Widom, 1998). Histone deposition and correct nucleosome folding on *λ* DNA were verified by electrophoretic mobility shift assay (EMSA) and micrococcal nuclease (MNase) protection assay (Figures 1C-F). Both assays were performed under native conditions allowing direct visualization of the fluorescent histone within intact nucleosomes and the DNA through SYBR Gold staining (Figures 1C and 1D). The deposition of histone octamers on *λ* DNA leads to size increase and is manifested in EMSA by the apparent shift of the *λ* DNA band (relative to naked *λ*), detected by both SYBR Gold and histone fluorescence (Figure 1C). The native MNase assay (Figures 1D and 1F), in which histone-free DNA is preferentially digested by MNase without subsequent deproteination, confirmed correct nucleosome formation. MNase digest of all tested nucleosomal *λ* templates produced intact mono- and di-nucleosomes whereas naked *λ* DNA was fully digested. The intercalating DNA stain SYBR Gold binds inefficiently to DNA wrapped around histone octamers resulting in a weaker DNA signal in the native MNase protection assay (and EMSA of saturated nucleosomal templates). Thus, in addition, we conducted a denaturing MNase protection assay (Figures 1E and 1F), in which the MNase-treated nucleosomal *λ* DNA templates were further subjected to proteinase K digest, yielding the expected histone-protected DNA fragments of ∼150 and ∼300 bp, consistent with mono- and di-nucleosome protected DNA, respectively. Taken together, these results clearly demonstrate the correct folding of WT and fluorescent nucleosomes on *λ* DNA.

### Fluorescent nucleosomes on λ DNA are discretely distributed in a ‘beads-on-a-string’ manner

By varying the octamer:DNA molar ratio in our reconstitution reactions we achieved different levels of nucleosome saturation (Figure 2). The deposition of increasing amounts of histone octamer on *λ* DNA was illustrated in EMSA by a steady increase in histone fluorescence and a gradual shift of the *λ* DNA band (Figures 2A and 2B). The associated native MNase protection assay revealed a corresponding increase in the amount of protected nucleosomal species (mono-, di- and tri-nucleosomes), confirming that the observed increase in the template size is due to correct nucleosome formation rather than non-specific histone-DNA interactions.

**Figure 2.**
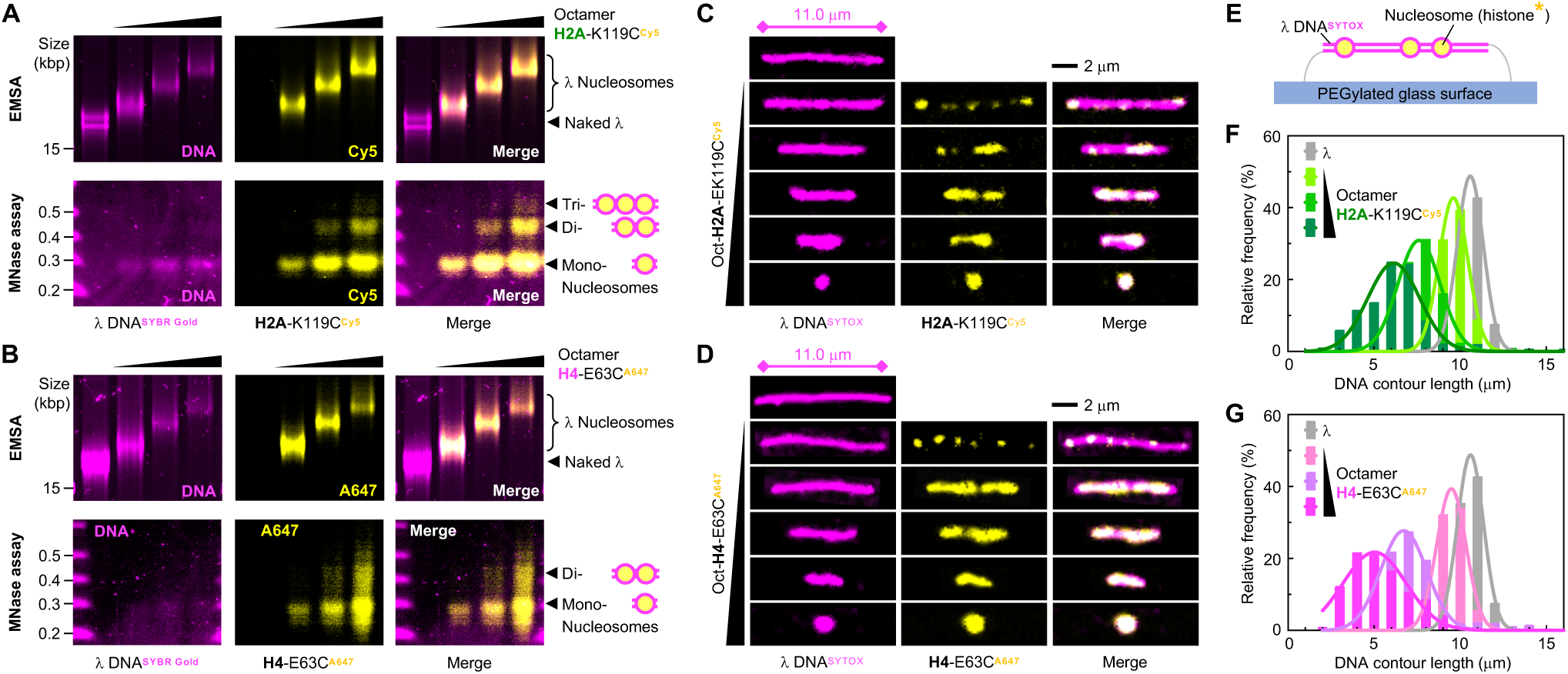
Fluorescent nucleosomes on λ DNA are discretely distributed in a ‘beads-on-a-string’ manner. (A and B) Native EMSA (top) and MNase assay (bottom) for nucleosomes labelled at H2A-K119C with Cy5 (A) and H4-E63C with AlexaFluor647 (B) reconstituted on *λ* DNA at increasing octamer:DNA ratios. Left panels show SYBR Gold staining of the DNA (magenta), central panels show Cy5 and AlexaFluor647 fluorescence signal (yellow) of labelled histones and right panels are the composites of both detection modes. Deposition of increasing amounts of histone octamer on *λ* DNA leads to gradual increase in the template size and slower migration through 0.5 % agarose in EMSA. The larger the template size the slower the migration, as manifested by the more prominent shift of the DNA band. The observed template size increase results from higher density of correctly folded nucleosomes as indicated by the presence of mono-, di- and tri-nucleosomes in the corresponding native MNase protection assays. The apparent loss of H4-E63C^A647^ signal in EMSA is most likely due to self-quenching of histone fluorescence, caused by structural arrangement of high-density nucleosomes. (C and D) Single-molecule imaging of nucleosomes labelled at H2A-K119C with Cy5 (C) and H4-E63C with AlexaFluor647 (D) reconstituted on *λ* DNA at increasing nucleosome density. Left panels show SYTOX Orange staining of the DNA (magenta), central panels show Cy5 and AlexaFluor647 fluorescence signal (yellow) of labelled histones and right panels are the composites of both detection modes. For details of experimental set up see panel E. Fluorescent nucleosomes reconstituted on *λ* DNA by salt dialysis show the characteristic ‘bead-on-a-string’ appearance. Nucleosome formation on *λ* DNA leads to apparent shortening of the DNA template, consistent with its wrapping around the octameric histone core. (E) Schematic of the DNA immobilized in the microfluidic device for single-molecule imaging. Fluorescent nucleosomes are pre-assembled on *λ* DNA by salt dialysis. The nucleosomal DNA template is stretched under flow and doubly tethered to the PEGylated glass surface of the microfluidic device via biotin-streptavidin interactions. The imaging is carried out in TIRF mode using 561- and 640-nm lasers to visualize SYTOX Orange-stained DNA (magenta) and Cy5/AlexaFluor647-labelled histones (yellow), respectively. (F and G) Single-molecule quantification of the DNA contour length for nucleosomes labelled at H2A-K119C with Cy5 (F) and H4-E63C with AlexaFluor647 (G) reconstituted on *λ* DNA at increasing octamer:DNA ratios. The four species presented on each graph correspond to the four samples shown in panels A and B. The DNA length of individual molecules was measured based on SYTOX Orange staining of the DNA (approximately 400 molecules at each histone octamer concentration). As illustrated in panels C and D, deposition of nucleosomes on *λ* DNA results in apparent shortening of the DNA template. The higher the octamer content in the reconstitution reaction, the shorter the mean DNA contour lengths and the broader the DNA length distributions were observed.

To visualize fluorescent nucleosomes on individual *λ* DNA molecules, we used microfluidics devices with a PEGylated and streptavidin-functionalized glass surface in combination with total internal reflection fluorescence (TIRF) microscopy (Figures 2C-G). Biotinylated *λ* DNA molecules containing pre-assembled fluorescently-labelled nucleosomes were first stretched under flow (to approximately 70% of the maximally stretched form) and tethered at both ends to the surface. The fluorescent DNA stain SYTOX Orange was then introduced into the chamber and both the DNA and fluorescent histone in nucleosomes were imaged in TIRF using 561- and 640-nm lasers, respectively (Figures 2C-E). Figures 2C and 2D show the images of individual *λ* molecules at increasing density of nucleosomes labelled at H2A-K119C with Cy5 and H4-E63C with AlexaFluor647, respectively. Fluorescent nucleosomes are discretely distributed on SYTOX-stained *λ* DNA as ‘beads-on-a-string’. As more nucleosomes are deposited on the DNA, the associated fluorescence signal of the labelled histone increases and appears more contiguous. Consistent with the DNA wrapping around the octameric histone core, we also observed apparent shortening of the *λ* DNA contour length upon nucleosome deposition, in a nucleosome density (histone octamer concentration)-dependent manner. We quantified this ‘shortening’ effect for samples shown in Figures 2A and 2B by measuring the contour length of approximately 400 individual molecules per sample and plotting the histograms (Figures 2F and 2G). As expected, the mean contour length decreased for nucleosomal samples reconstituted with higher concentrations of histone octamer and the length distributions became broader. At very high histone octamer:DNA ratios, the molecules appeared as intense diffraction-limited spots of fluorescence (Figures 2C and 2D, bottom panels) that, in contrast to singly-tethered low density nucleosomal templates, did not stretch under buffer flow (Videos 1 and 2).

### Assay for single-molecule imaging of parental histones during DNA replication

To investigate the dynamics of parental histones during DNA replication, we combined real-time TIRF imaging with microfluidics-based replication assays in nucleus-free *Xenopus laevis* egg extracts (Figure 3A) (Lebofsky et al., 2009; Loveland et al., 2012; Yardimci et al., 2012). Stretched *λ* DNA containing pre-assembled fluorescent nucleosomes (one of the histones labelled fluorescently with either Cy5 or AlexaFluor647) was attached to the surface of the microfluidic flow cell as described above. The nucleosomal templates were then incubated for approximately 15 minutes in a high-speed supernatant (HSS) of *Xenopus* eggs to promote sequence non-specific origin licensing (i.e. the ORC-dependent assembly of pre-replication complexes). Next, a concentrated nucleoplasmic extract (NPE) was introduced into the microfluidic chamber, which initiates and supports efficient bidirectional replication. The number of replication initiations per DNA template was regulated by adding the Cdk2 inhibitor p27^Kip^ (Walter and Newport, 2000; Yardimci et al., 2010). To allow visualization of replication fork progression in real time, NPE was supplemented with a fluorescent fusion protein Fen1-KikGR, which decorates replication bubbles but does not detectably alter ensemble replication kinetics or replication bubble growth monitored in single-molecule assays. Fen1-KikGR fluorescence was detected using 488-nm laser (Loveland et al., 2012).

**Figure 3.**
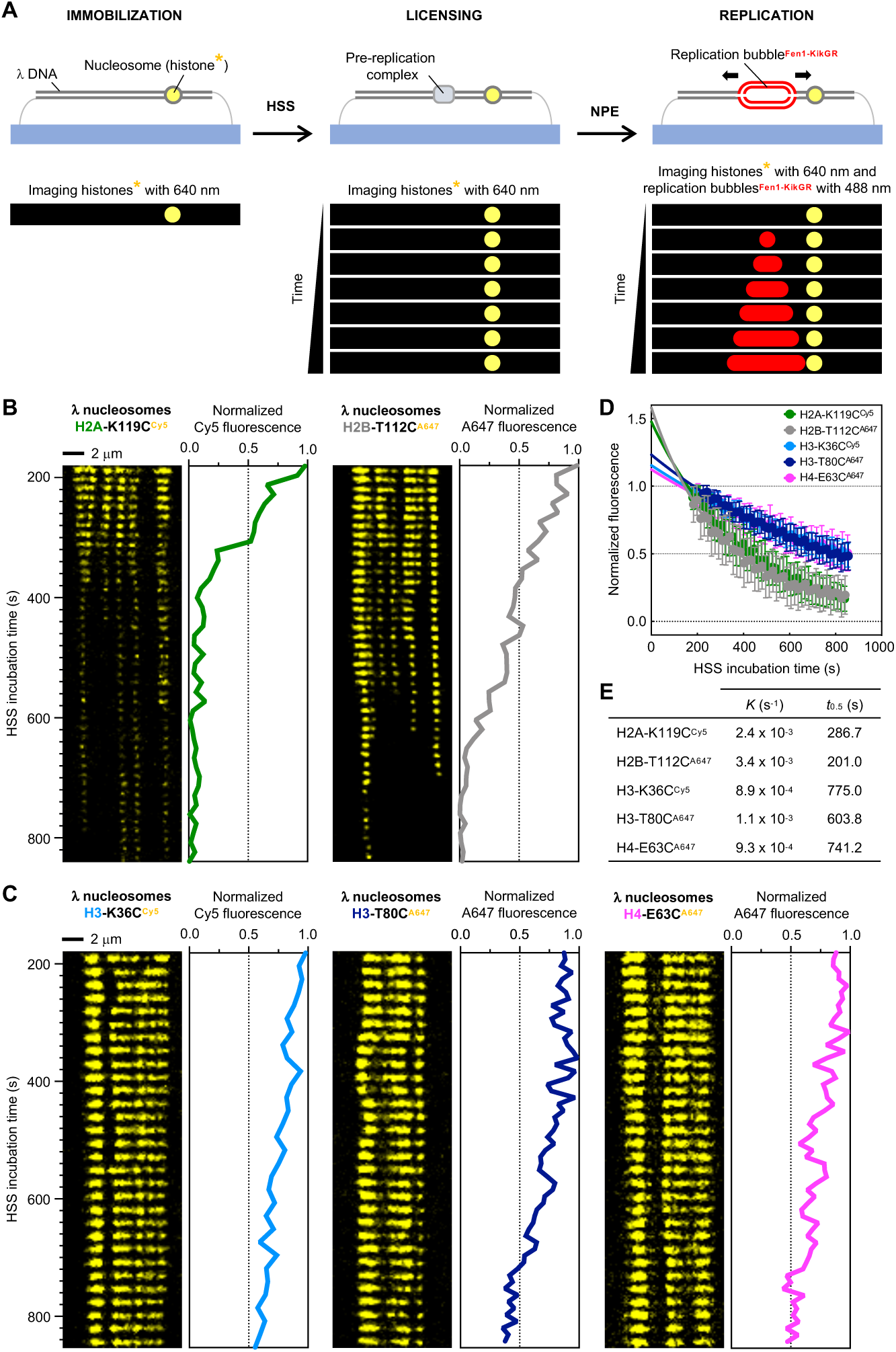
Histone dynamics during DNA licensing in HSS. (A) Schematic of the experimental set-up for real-time single-molecule imaging of nucleosome dynamics during replication in *Xenopus leavis* egg extracts. *λ* DNA containing fluorescent nucleosomes (one of the four histones labelled fluorescently) is stretched under flow and tethered at both ends to the functionalized glass surface of a microfluidic flow cell. The immobilized DNA is licensed in high-speed supernatant (HSS). Bidirectional replication is initiated upon introduction of nucleoplasmic extract (NPE) supplemented with a fluorescent fusion protein Fen1-KikGR, which decorates replication bubbles and allows progression of replication forks to be tracked in real time. Cy5- or Alexa647-labelled histones within immobilized nucleosomal templates are imaged with a 640-nm laser at each stage. Replication fork progression is visualized in NPE using a 488-nm laser. (B and C) Kymograms and corresponding intensity profiles for fluorescent *λ* nucleosomes during incubation in HSS. Nucleosomes labelled at H2A-K119C with Cy5 and H2B-T112C with AlexaFluor647 (B) show faster loss of fluorescence than nucleosomes labelled at H3-K36C with Cy5, H3-T80C with AlexaFluor647 and H4-E63C with AlexaFluor647 (C). (D) Plot showing the mean loss of fluorescent signal for *λ* nucleosomes (H2A-K119^Cy5^, H2B-T112C^A647^, H3-K36C^Cy5^, H3-T80C^A647^ and H4-E63C^A647^) during incubation in HSS. Over 100 molecules were analyzed for each histone template. Individual fluorescence decay traces were normalized to background (‘0’) and maximum value of fluorescence (‘1’). A mean fluorescence value and standard deviation were calculated and plotted for each time point. The mean value traces were then fitted to a single exponential function. (E) Summary of the fluorescence decay rate constants (*K*) and half-lives (*t*0.5) extracted from the single exponential fit to the data presented in panel C. See Table S2 for detailed fitting parameters.

### Histone dynamics during DNA licensing – Differential exchange kinetics of H2A/H2B versus H3/H4

We first investigated histone dynamics during DNA licensing (Figure 3 and Videos 3–7). HSS was introduced into the flow cell containing immobilized fluorescently-labelled *λ* nucleosomes at a low flow rate over 2.5 minutes. As the extract slowly reached *λ* nucleosomes, thermal fluctuations of individual molecules become gradually reduced in comparison to egg lysis buffer (ELB) conditions. We rationalized that this is due to the immobilized DNA being bound by extract proteins, including native histones. Further chromatinization in extracts also brought stretched *λ* DNA molecules closer to the surface resulting in an initial increase in the histone fluorescence intensity (Figure S1A). Thus, to obtain reliable data on histone dynamics in HSS, we imaged histones in real time between 3 and 14 min of incubation, under no flow conditions and after the initial changes in molecule mobility and fluorescence intensity.

Figures 3B and 3C show kymograms and corresponding fluorescence traces for *λ* nucleosomes containing H2A-K119^Cy5^, H2B-T112C^A647^, H3-K36C^Cy5^, H3-T80C^A647^ and H4-E63C^A647^. We found that all analyzed fluorescent histones show limited lateral dynamics; i.e. their movement along the DNA molecule is largely confined within the spatiotemporal resolution of our approach. While there was some local drift of histone fluorescence from the starting position, we never observed long-distance lateral movement for any of the histones tested in our assay. The dominant dynamic behavior observed during the HSS incubation is the gradual loss of histone-associated fluorescence over time. Interestingly, in the case of H2A-K119^Cy5^ and H2B-T112C^A647^ (Figure 3B, Videos 3 and 4) the decrease in histone fluorescence signal was greater than for H3-K36C^Cy5^, H3-T80C^A647^ and H4-E63C^A647^ (Figure 3C, Videos 5, 6 and 7). We quantified the average rate of histone fluorescence decay for each template (Figures 3D, 3E and S1B, and Table S2) and found that histones H3 and H4 have approximately three-times longer half-lives in HSS than H2A and H2B.

The observed loss of histone fluorescence is likely to result from three phenomena: (i) photobleaching of the dye, (ii) nucleosome eviction and (iii) fluorescent histone exchange within a nucleosome with an unlabelled native counterpart from the extract. Given that the same type of dye was used to track histones displaying different kinetic behavior (e.g. H2B-T112C^A647^ and H4-E63C^A647^; half-lives: 201.0 and 741.2 s^-1^, respectively) and that histone H3 labelled with two different dyes showed similar rate of fluorescence decay (H3-K36C^Cy5^ and H3-T80C^A647^; half-lives: 775.0 and 603.8 s^-1^, respectively), we conclude that photobleaching itself cannot account for the observed kinetic differences between H2A/H2B and H3/H4. We also rationalized that nucleosome eviction would affect the fluorescence signal in the same way, regardless of the histone type, as all four histones would simultaneously dissociate from the DNA. Hence, we conclude that the observed difference in the loss of fluorescence between H2A/H2B and H3/H4 predominantly reflects different exchange rates with native histones, present in HSS at a concentration of ∼1-6 μM (Figure S1C). The faster displacement rate for histones H2A and H2B relative to H3 and H4 in *Xenopus* extracts is consistent with previous reports indicating greater lability of H2A-H2B within nucleosomes *in vivo* (Annunziato, 2013; Jackson, 1990; Kimura and Cook, 2001; Louters and Chalkley, 1985) and could potentially reflect the structural organization of the histone octamer, where the two H2A-H2B dimers are more accessible than the core (H3–H4)_2_ hetero-tetramer (Figure 1A).

### Replication of fluorescent nucleosomal templates

To investigate the dynamics of parental histones during DNA replication, we initiated replication of the stretched and licensed fluorescent nucleosomal *λ* templates by introducing NPE containing Fen1-KikGR (Figure 3A). After one or two origins per template had fired, the NPE mix was replaced with NPE supplemented with p27^Kip^ to prevent further origin firing. This procedure allowed us to follow the growth of individual replication bubbles in real time, and hence to determine the outcome of collision between a single progressing replication fork with nucleosomes on its path. We anticipated that as long as fluorescent nucleosomes are sparsely distributed along the stretched DNA molecules (i.e. a few fluorescent nucleosomes per DNA molecule), we would be able to distinguish individual fork-nucleosome collision events.

We first examined whether replication of nucleosomal *λ* templates is as efficient as that of naked *λ*. To this end, we measured the mean replication fork velocity for naked *λ*, as well as WT and fluorescent nucleosomes on *λ* DNA. For all templates, the replication forks travelled at a similar mean velocity of approximately 640 nt/min (Figure S2A and S2B), indicating that neither the reconstituted nucleosomes nor the fluorophores they carry affect fork progression. Forks fired on similar time scales for naked DNA and low density nucleosomal templates; between 5 and 12 min from the moment of NPE introduction. In addition, we compared the efficiency of replication using ensemble assays, in which naked plasmid and plasmid containing fluorescent nucleosomes were replicated under unrestricted firing conditions in *Xenopus* egg extracts. We found that chromatinized plasmids replicated as efficiently as their naked counterpart (Figure S2C). We conclude that our nucleosomal *λ* templates and microfluidics-based replication assays provide an appropriate imaging platform for tracking the fate of parental histones during replication.

### Heterogenous dynamics of parental histones upon replication fork arrival

The primary goal of this study was to determine the outcome of replication fork encounters with parental nucleosomes by using our real-time single-molecule imaging platform. Based on the literature, we envisioned four possible collision scenarios, all of which would give us a characteristic kymogram footprint (Figure 4; schematics). These included nucleosome (histone) eviction, localized parental histone transfer onto daughter strands, nucleosome (histone) sliding ahead of the replication fork and replication fork stalling. We focused our studies on low nucleosome density *λ* templates containing either H3-K36C^Cy5^ or H4-E63C^A647^, due to their high fluorophore labelling efficiency and lower exchange rates, compared to H2A-H2B, during licensing in HSS. We observed all of these events in our experiments (Figures 4 and S3).

**Figure 4.**
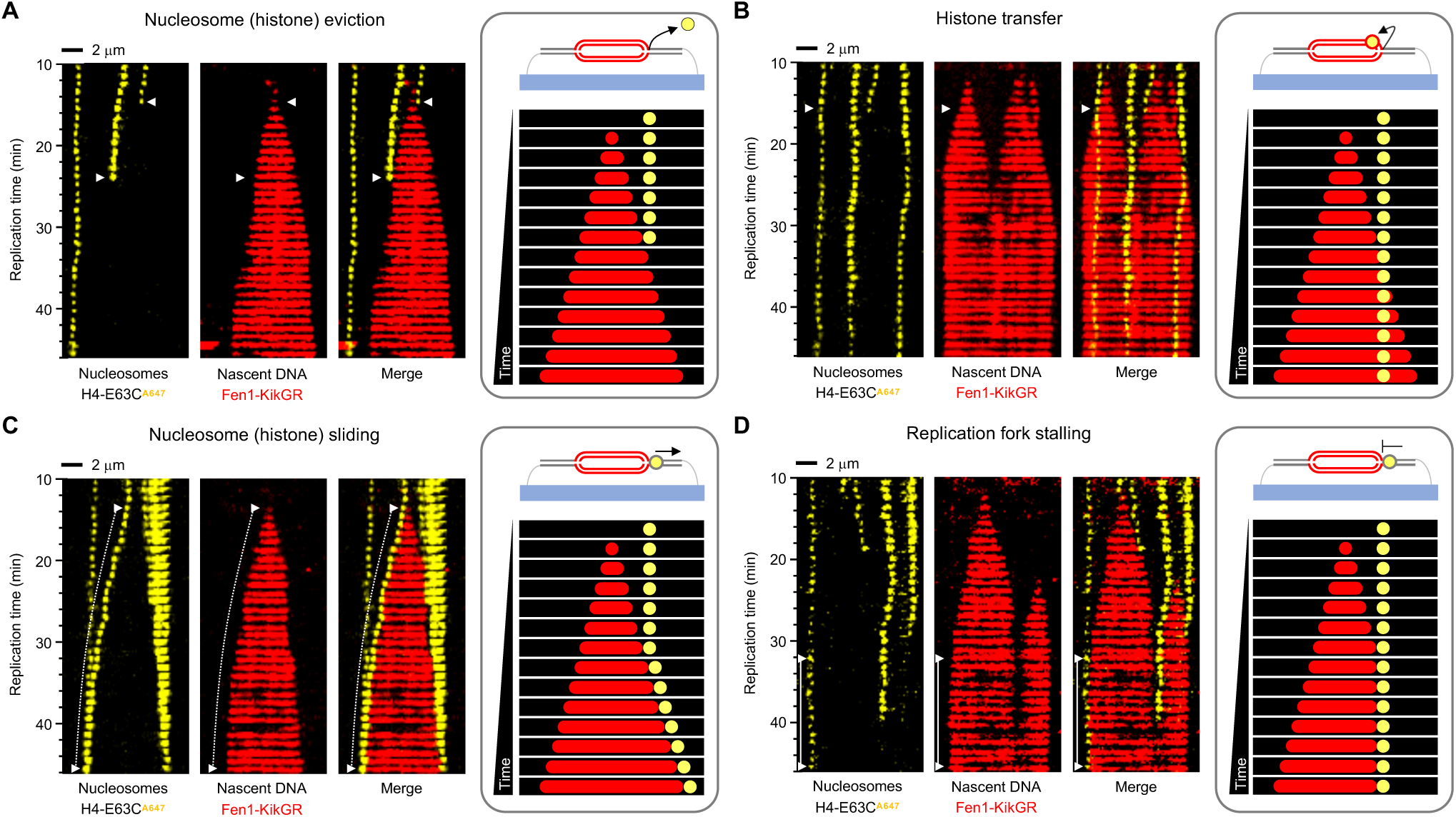
Heterogenous dynamics of parental histones upon replication fork arrival. For each specified outcome, data are presented as kymograms of nucleosome-associated fluorescence (H4-E63C^A647^; yellow; left panels), Fen1-KikGR signal indicating nascent DNA (red; central panels) and both signals together (merge; right panels). Time and size scales are presented. The white triangles mark the point of initial encounter between the replication fork and nucleosome. Dotted lines indicate sliding events, whereas solid lines correspond to replication fork stalling. For clarity, a schematic representation of each outcome is shown in grey borders. (A) Nucleosome (histone) eviction is manifested by the loss of histone fluorescence at the point of collision with the replication fork. It represents the disassembly of nucleosomes ahead of the replication fork. (B) Histone transfer is observed when the histone-associated fluorescence is retained and incorporated into the track of replicated DNA. This event illustrates successful localized parental histone recycling, whereby a histone released from parental DNA ahead of the replication fork is kept in the vicinity of the replisome, followed by its deposition onto daughter DNA. (C) Nucleosome (histone) sliding is observed when the histone-associated fluorescence moves together with the tip of the replication bubble (marked as a dotted white line). This behavior is consistent with two molecular phenomena; either the whole nucleosome is pushed ahead of the replication fork or a parental histone released from the DNA associates with the replisome and travels with it along the DNA. (D) Replication fork stalling occurs when nucleosome constitutes a roadblock preventing the replication fork from further movement. It is manifested in the kymogram as an arrested tip of the replication bubble next to a static histone signal (indicated as a solid line).

Nucleosome (histone) eviction is emblematic of nucleosome disassembly prior to DNA unwinding and synthesis, resulting in parental histone release into the pool of free histones. It is manifested in the kymograms and accompanying videos by the loss of histone fluorescence at the point of encounter with the replication fork (Figure 4A, Video 8 and Figure S3A). In the case of nucleosome (histone) transfer, the histone-associated fluorescence is incorporated into the Fen1-KikGR-decorated track of nascent DNA upon passage of the replication fork (Figure 4B, Video 9 and Figure S3B). This characteristic illustrates localized parental histone recycling, a mechanism whereby the fluorescent histone from disassembled parental nucleosome stays in the vicinity of the replisome and is immediately re-deposited into a nucleosome on daughter DNA. The resolution of our technique is approximately 1 kilobase pair, and so it does not allow us to specify if histones are reinstated at the exact same locus within the replicated DNA. The third type of event, which we classify as nucleosome or histone sliding, is detected as a continuous movement of histone fluorescence signal with a tip of the replication bubble from the moment of nucleosome-fork encounter (Figure 4C, Video 10 and Figure S3C). This sliding behavior is likely indicative of two molecular phenomena, which at present cannot be distinguished. One possibility is that the whole nucleosome is being pushed ahead of the replication fork, as observed for nucleosome remodelers (Bowman, 2010). Alternatively, the nucleosome is disassembled at the point of fork collision, the fluorescent histone then associates with the replisome and travels with it along the DNA. Sliding typically occurs over short distances (within a few kilobase pairs) but occasionally we observed nucleosome/histone push on a scale of 25 – 30 kilobase pairs, spanning over a half of the length of *λ* DNA (48.5 kbp). Replication fork stalling upon collision with a nucleosome is exemplified in our experiments by a static histone fluorescence next to an arrested tip of the replication bubble (Figure 4D, Video 11 and Figure S3D). In this scenario, the nucleosome acts as a roadblock preventing the replication fork from further movement.

Nucleosome eviction and localized histone transfer are the two ultimate outcomes of replication fork encounter with nucleosomes as, once they have occurred, the fork and nucleosome (histone) are no longer in contact/proximity. In contrast, nucleosome/histone sliding and replication fork stalling preserve the fork-nucleosome/histone ‘interaction’, and hence often lead to secondary outcomes (Figure 5). Both sliding and stalling can terminate in nucleosome/histone eviction (Figures 5A and 5B, and Videos 12 and 13, respectively) as well as histone transfer behind the replication fork (Figures 5C and 5D, and Videos 14 and 15, respectively). In addition, nucleosome/histone sliding can result in replication fork stalling (Figure 5E and Video 16) and vice versa (Figure 5F and Video 17). Occasionally, we observe tertiary events; for example, fork stalling followed by nucleosome/histone sliding leads to a second fork stalling (Figure 5F, note that the second fork stalling event is unmarked) or nucleosome/histone sliding followed by fork stalling terminates in histone transfer (Figure S6D; note that fork stalling and histone transfer are unmarked).

**Figure 5.**
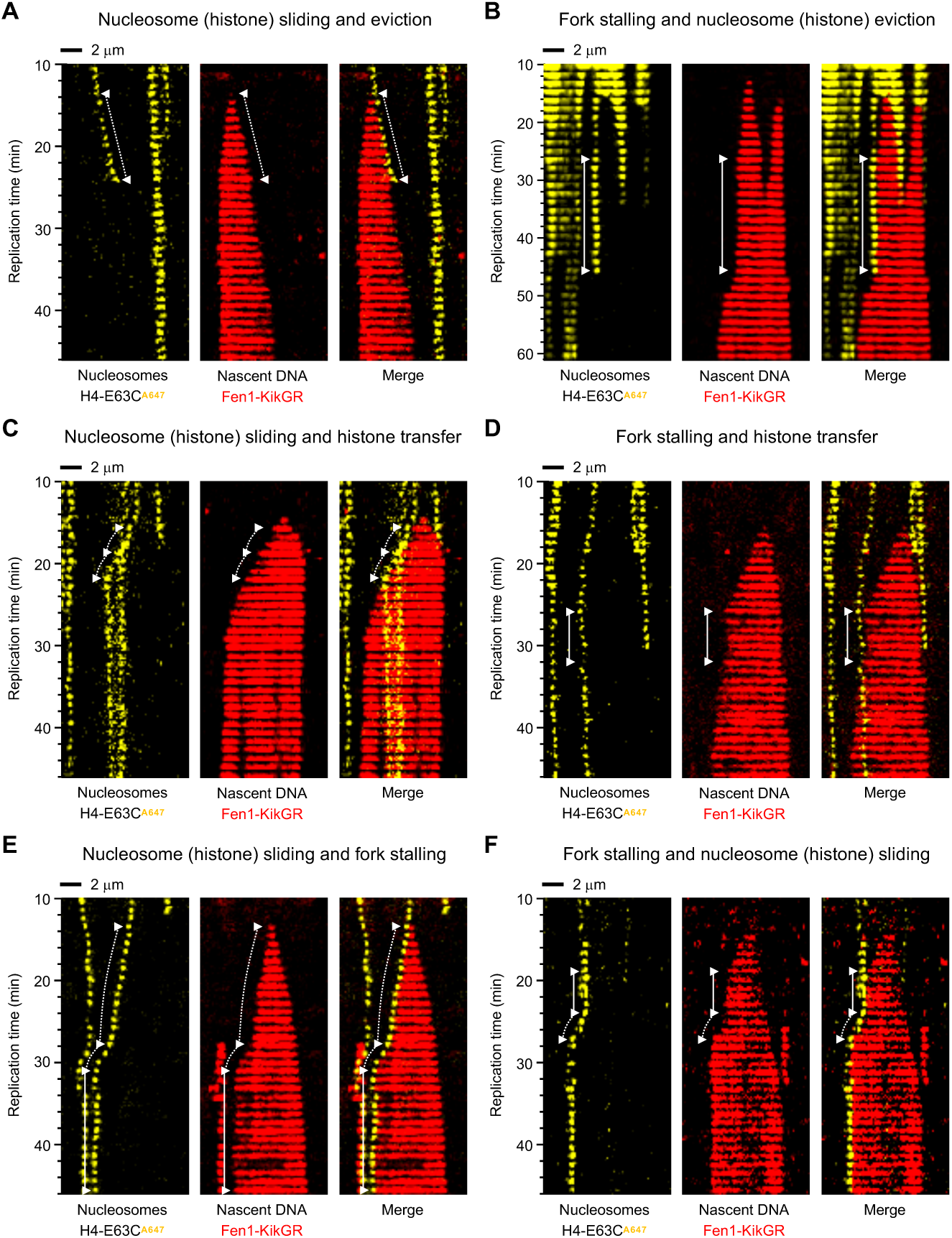
Secondary outcomes of the replication fork collision with nucleosomes during DNA replication in *Xenopus* egg extracts. For each specified outcome, data are presented as kymograms of nucleosome-associated fluorescence (yellow; left panels), Fen1-KikGR signal indicating nascent DNA (red; central panels) and both signals together (merge; right panels). Time and size scales are presented. The white triangles mark the point of initial encounter between the replication fork and nucleosome. Dotted lines indicate sliding events, whereas solid lines correspond to replication fork stalling. (A, C and E) Nucleosome (histone) sliding can terminate in eviction (A), histone transfer (C) and replication fork stalling (E). (B, D and F) Replication fork stalling can lead to nucleosome (histone) eviction (B), histone transfer (D) and nucleosome (histone) sliding (F).

### Parental histone eviction is the dominant outcome of replication fork encounter with nucleosomes in *Xenopus* egg extracts

To gain further insight into the mechanism of chromatin replication, we quantified the probability of different outcomes of fork-nucleosome encounter in *Xenopus* egg extracts (Figures 6A and 6B; left panels). Contrary to our expectations, for both tested nucleosomal templates, containing either H4-E63C^A647^ or H3-K36C^Cy5^, the dominant event was nucleosome eviction at 40.2% and 49.1%, respectively. Parental histone recycling, the event we anticipated to be the most frequent, occurred at significantly lower frequency, 15.4% for H4-E63C^A647^ nucleosomes (the rarest event of all) and 17.3% in the case of nucleosomes carrying H3-K36C^Cy5^. Nucleosome/histone sliding was more prevalent on templates with H4-E63C^A647^ nucleosomes (27.4%) than H3-K36C^Cy5^ (19.1%) however, in both cases, it represented the second most probable outcome of replication fork collision with nucleosomes. Replication forks stalled on nucleosomes in 16.9% of cases for H4-E63C^A647^ and 14.5% for H3-K36C^Cy5^.

**Figure 6.**
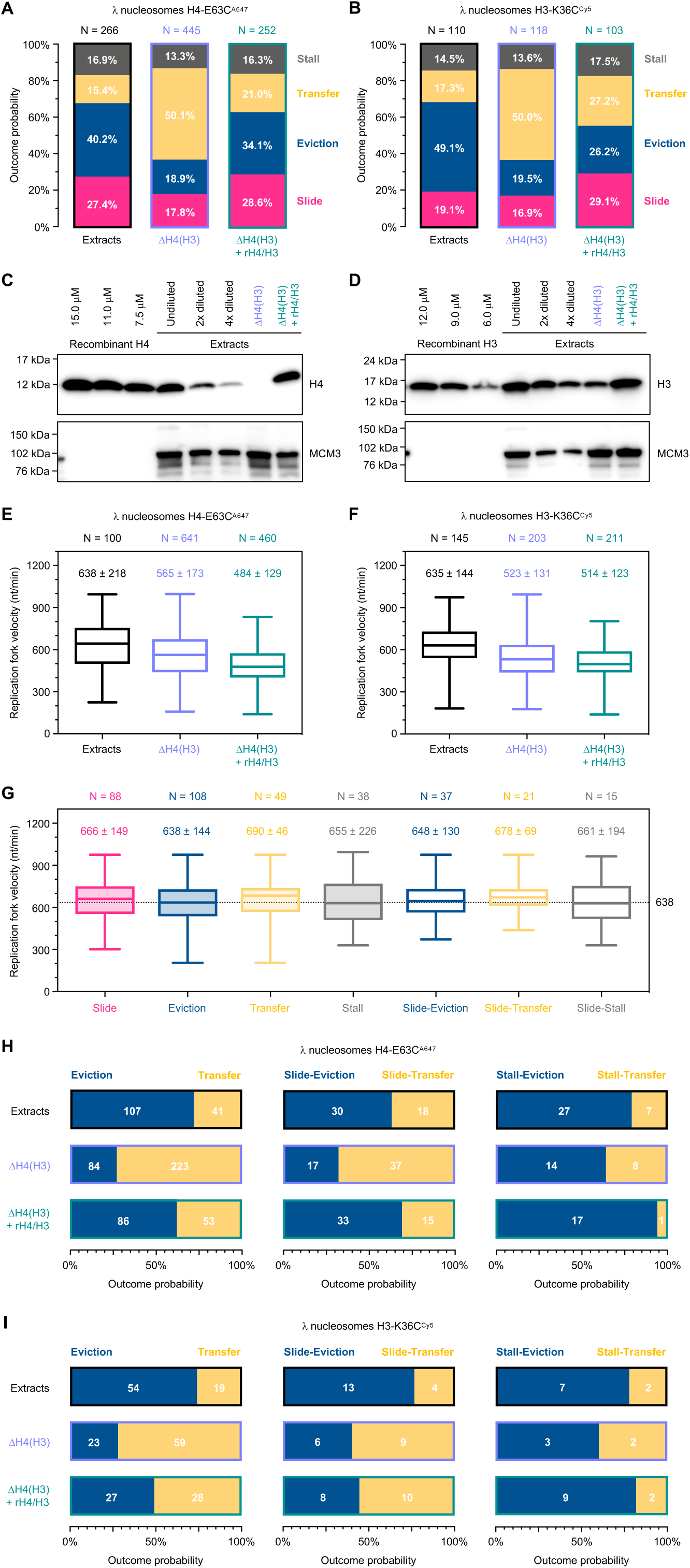
Effect of free histones on parental histone dynamics at the replication fork. (A and B) Quantification of the four basic outcomes of replication fork collision with nucleosomes labelled at H4-E63C^A647^ (A) and H3-K36C^Cy5^ (B) in regular extracts (black borders; left panels; a mix of HSS and NPE at 1:1 volume ratio), extracts depleted of histone H4 and H3 (ΔH4/H3; blue borders; central panels) and depleted extracts supplemented with recombinant histones H3 and H4 (ΔH4/H3 + rH4/H3; green borders; right panels). Nucleosome/histone eviction is shown in blue, histone transfer in yellow, nucleosome/histone sliding in pink and replication fork stalling in grey. N indicates the total number of analyzed collisions and contributions from different outcomes are specified. Data from at least two biological repeats were pooled in the analysis for each tested condition. (C and D) Western blots used to estimate the concentration of histone H4 (C) and H3 (D) in regular extracts, extracts depleted of histone H4 and H3 (ΔH4/H3) and depleted extracts supplemented with recombinant histones H3 and H4 (ΔH4/H3 + rH4/H3). (E and F) Box-and-whisker Tukey plot of replication fork velocities measured in regular extracts (black), extracts depleted of histone H4 and H3 (ΔH4/H3; blue) and depleted extracts supplemented with recombinant histones H3 and H4 (ΔH4/H3 + rH4/H3; green) for *λ* nucleosomes containing H4-E63C^A647^ (E) and H3-K36C^Cy5^ (F). Values above the box plots indicate the mean replication fork velocity extracted from the Gaussian fit, plus and minus standard deviation. The number of values analyzed per data set (N) is also shown. (G) Box-and-whisker Tukey plot of replication fork velocities measured in regular extracts and categorized by the collision outcome. Data for *λ* nucleosomes containing H4-E63C^A647^ and H3-K36C^Cy5^ were pooled to generate this plot. Values above the box plots indicate the mean replication fork velocity extracted from the Gaussian fit, plus and minus standard deviation. The number of values analyzed per data set (N) is also shown. The horizontal dotted line marks the mean replication fork velocity in regular extracts, 638 nt/min. (H and I) Quantification of nucleosome/histone eviction (blue) versus histone transfer (yellow) for nucleosomes labelled at H4-E63C^A647^ (H) and H3-K36C^Cy5^ (I). Analysis for primary (eviction versus transfer) and secondary (slide/stall-eviction versus slide/stall-transfer) outcomes is presented.

In the light of this surprisingly inefficient parental histone recycling at the replication fork, we next wanted to test whether DNA stretching could influence the outcome of nucleosome-fork collision. It is plausible that, if a specific three-dimensional fork-replisome structure is needed for efficient parental histone transfer onto daughter strands, the double-tethering of nucleosomal templates (stretched to ∼70% of the maximum contour length) could potentially impede longer-range DNA contacts and histone recycling. To address this issue, we performed single-molecule replication experiments, in which approximately 50% of the immobilized nucleosomal *λ* molecules were tethered to the surface at only one end, and so were free to fold in three-dimensional space, unlike their doubly-tethered counterparts (Figure S4). Singly-tethered molecules in extracts appear as a diffraction-limited spot of fluorescence, which does not stretch under flow of native buffers, making it impossible to visualize individual fork collisions with nucleosomes. Hence, we compared the loss of histone-associated fluorescence between singly- and doubly-tethered nucleosomal templates as a proxy for determining the effect of DNA stretching on parental histone retention during replication. We found no difference in the rate of histone loss or daughter strand synthesis (as measured by the increase of the Fen1-KikGR signal) between the two templates (Figures S4D and S4E). Based on these observations we conclude that DNA stretching does not cause excessive histone eviction during replication in *Xenopus* egg extracts.

Another potential cause of inefficient histone recycling at the replication fork in our system could be the retention of Fen1-KikGR on nascent DNA. To test whether this is the case, we conducted single-molecule replication experiments on doubly-tethered fluorescent *λ* nucleosomes in extracts supplemented with digoxygenin-dUTP (dig-dUTP), instead of Fen1-KikGR (Figure S5). Incorporation of dig-dUTP into nascent DNA does not allow us to track the growth of replication bubbles in real time but it enables their post-replication visualization through immunostaining with fluorescein-labelled anti-digoxygenin antibody (anti-dig Ab^Fluor^). We combined three modes of detection after replication (nucleosomes – H3-K36C^Cy5^, nascent DNA – anti-dig Ab^Fluor^, all DNA – SYTOX Orange) and found that the replicated tracts of *λ* DNA were largely free of H3-K36C^Cy5^ signal, whereas the non-replicated *λ* regions remained decorated with H3-K36C^Cy5^-nucleosomes (Figure S5B). These results further confirm that histone recycling is highly inefficient during replication in *Xenopus* egg extracts and lead us to conclude that Fen1-KikGR does not interfere with histone transfer onto daughter strands.

### Efficiency of parental histone transfer depends on the concentration of free histones

In *Xenopus leavis* embryos, transcription is activated in the thirteenth cell cycle (Newport and Kirschner, 1982). Until this point, the embryonic genome is transcriptionally silent, and so the oocyte must provide histones in sufficient abundance to support the initial twelve rounds of replication after fertilization (Woodland et al., 1979). *Xenopus* egg extracts must therefore contain a high proportion of free histones; at least 2^12^ times higher than an equivalent extract of somatic cells. Thus, we set out to determine whether the probabilities of the four outcomes of fork-nucleosome encounter would be different in extracts containing less histones.

We estimated the concentration of histones in our replication-promoting extracts by Western blots as approximately 10 and 20 μM for H4 and H3, respectively (Figures 6C and 6D). Newly synthesized histone H4 is acetylated at lysine 12 (H4-K12Ac) and forms a pre-deposition complex with histone H3 (Verreault et al., 1998). We depleted extracts of histone H4, using an antibody recognising H4-K12Ac (Zierhut et al., 2014), to less than 10% of its normal content; estimated concentration of H4 in depleted extracts is ∼1 μM (Figure 6C). This procedure also led to co-depletion of histone H3 from extracts and reduced its concentration to ∼5 μM (equivalent to 25% of its normal content; Figure 6D). We next performed single-molecule replication assays on doubly-tethered *λ* nucleosomes, containing either H4-E63C^A647^ or H3-K36C^Cy5^, in extracts depleted of histones H4 and H3. For both templates, we observed a reduction in the mean replication fork velocity relative to regular extracts (565 nt/min from 638 nt/min for H4-E63C^A647^ and 523 nt/min from 635 nt/min for H3-K36C^Cy5^; Figures 6E and 6F). Based on the observation that H4/H3-depleted extracts contain less histone chaperone Asf1 (∼25% less than in undepleted extracts; Figure S6A), we rationalized that the observed reduction in the replication fork rates might reflect changes in the histone-to-chaperone ratio. The four principal outcomes of fork-nucleosome encounter were still detected in depleted extracts (Figures S6B-E) but the probability of collision outcomes was different (Figures 6A and 6B; central panels), in particular, regarding parental histone transfer and nucleosome eviction (Figures 6H and 6I). In stark contrast to regular extracts, the dominant event in histone-depleted extracts was localized histone transfer, detected in 50% of collisions for both H4-E63C^A647^ and H3-K36C^Cy5^ *λ* nucleosomes. This increased efficiency of histone recycling was accompanied by a dramatic drop in the frequency of nucleosome/histone eviction (18.9% for H4-E63C^A647^ and 19.5% for H3-K36C^Cy5^), whereas nucleosome/histone sliding and replication fork stalling were observed at similar probability levels to those found in undepleted extracts. We also observed a higher probability of secondary transfer events (i.e. slide-transfer and stall-transfer), when compared to regular extracts (Figures 6H, 6I and S7).

Given the lower mean replication fork velocity in extracts depleted of histones H4 and H3, we next investigated whether the observed increase in localized histone transfer is due to slower replication forks. If that was the case, in regular undepleted extracts, the mean velocity of forks leading to histone transfer upon collision with nucleosomes would be lower than for forks prompting nucleosome eviction. We compared replication fork velocities leading to different outcomes upon nucleosome-fork encounter in regular extracts and detected no such difference (Figure 6G). Indeed, we found no correlation between replication fork speed and any of the nucleosomal outcomes evident during replication in extracts.

Our results strongly suggest that excess provision of free histones during replication, as found in *Xenopus* egg extracts, leads to impaired localized histone recycling. We further tested this hypothesis by performing single-molecule replication assays in extracts depleted of the free endogenous histones (as described above) but supplemented with recombinant histones H4 and H3 to native concentrations (Figures 6C and 6D). If our model is correct, the presence of recombinant histones should counteract the H4(H3) depletion effect and mimic the behavior of regular undepleted extracts. We replicated nucleosomal templates containing either H4-E63C^A647^ or H3-K36C^Cy5^ and detected a slight reduction in the mean replication fork velocity, in comparison to depleted extracts (484 nt/min from 565 nt/min for H4-E63C^A647^ and 514 nt/min from 523 nt/min for H3-K36C^Cy5^; Figures 6E and 6F). Next, we quantified the probability of different fork-nucleosome encounter outcomes in depleted extracts supplemented with recombinant histones (Figures 6A and 6B; right panels; and Figure S8). Consistent with our predictions, we found reduced levels of histone transfer (21.0% for H4-E63C^A647^ and 27.2% for H3-K36C^Cy5^) and higher frequency nucleosome/histone eviction events (34.1% for H4-E63C^A647^ and 26.2% for H3-K36C^Cy5^), relative to histone depleted extracts (Figures 6A, 6B, 6H and 6I). A similar trend was also observed for secondary transfer and eviction events (Figures 6H, 6I and S7); i.e. events following initial slide and stall. In the case of H4-E63C^A647^ *λ* nucleosomes, nucleosome/histone sliding and replication fork stalling were detected at similar probability levels to those found in regular and undepleted extracts (Figures 6A and 6H). We note that for *λ* nucleosomes containing H3-K36C^Cy5^ (Figures 6B and 6I) these two events were found at a slightly higher frequency than previously detected for regular and undepleted extracts. Based on these data, we conclude that the efficiency of localized histone recycling at the replication fork depends on the concentration of soluble histones.

## DISCUSSION

Chromatin domains and their constituent histones with specific PTMs define the transcriptional program of the cell, and hence must be faithfully replicated through cell division. During replication chromatin undergoes a complete nucleosome-by-nucleosome disassembly, followed by restoration of chromatin structures on the daughter strands. Due to the dynamic and multi-component nature of chromatin replication, the molecular mechanisms that govern nucleosome disassembly and parental histone transfer remain poorly characterized. In this work, we devised a real-time single-molecule imaging platform to determine the fate of parental nucleosomes and their constituent histones upon encounter with progressing replication forks. Our approach enables visualization of individual nucleosome-fork collisions during replication in *Xenopus* egg extracts, and thus allowed us to unravel the mechanistic detail of chromatin replication at an unprecedented spatiotemporal resolution. Broader implications and significance of our findings are discussed below.

### Implications of heterogenous parental histone dynamics upon collision with the replication fork

The current consensus model for replication-coupled parental histone transfer suggests that (i) most if not all parental histones are recycled at the replication fork (Alabert et al., 2015), (ii) parental histones are quickly deposited onto nascent DNA and are equally distributed between the leading and lagging strands (Annunziato, 2013, 2015; Petryk et al., 2018; Yu et al., 2018), (iii) parental histones are recycled with their specific PTMs (Alabert et al., 2015), and (iv) that genomic localization of parental histones is preserved on daughter strands (Reveron-Gomez et al., 2018). Most of these pioneering studies are based on tailored ChIP-Seq and proteomics approaches that, while yielding important insights into replication-coupled chromatin restoration in bulk, inevitably, average out any inhomogeneities. They also do not provide crucial information on time-resolved parental histone dynamics, since they compare only pre- and post-replicated states of chromatin. Our single-molecule nucleosome imaging methodology offers a complimentary tool to study chromatin dynamics, as it focuses on individual parental histones, tracks them through the entire process of DNA replication in real time, and thus unravels their intricate kinetic behavior.

Through the use of this approach, we managed to demonstrate that, contrary to the prevailing view, replication fork collision with nucleosomes does not always result in an instant parental histone transfer onto daughter strand (Figures 4 and 5; we note that at present our experiments do not allow us to determine whether a physical collision actually takes place between the replisome and nucleosomes). In fact, three additional outcomes are possible: nucleosome/histone eviction, nucleosome/histone sliding and replication fork stalling. While nucleosome/histone eviction undoubtedly represents parental nucleosome disassembly, the very first step on the possible histone recycling trajectory, the latter two cases have not been observed before for nucleosome-fork encounter. Nucleosome/parental histone sliding has two equally probable molecular explanations that, as yet, we cannot distinguish; either a whole nucleosome is pushed ahead of the replication fork or an evicted parental histone is ‘piggybacking’ on the replisome. The ‘piggybacking’ mechanism is particularly interesting since, if true, it would represent the second intermediate step on the histone transfer pathway, whereby released parental histones are ushered to daughter strands via a series of interactions facilitated by histone chaperones and replisome components, such as FACT, MCM2, Ctf4 or Pol*α* (Gan et al., 2018; Kurat et al., 2017). Further studies are needed to identify the underlying molecular basis for the observed sliding behavior.

Replication fork stalling upon collision with a nucleosome has an obvious molecular interpretation – a nucleosome constitutes a roadblock and stops progression of the replisome. Indeed, other DNA-binding proteins can lead to fork stalling in egg extracts (Kose et al., 2019). Fork stalling is a transient state that, in most cases, terminates in nucleosome eviction or parental histone recycling. Persistent stalling events (i.e. when replication fork never restarts on the experimental time scale) typically occur when the nucleosome is located at the very end of *λ* DNA. Because the DNA molecules in our assays are of finite length (48.5 kbp), the likelihood of finding an end-point nucleosome is much higher than for longer DNA, and so the proportion of persistent stalling events in our quantifications must be an overestimate. In addition, at present, we cannot draw any definitive conclusions as to what makes some nucleosomes more difficult to disassemble ahead of the replication fork particularly since the molecular basis for nucleosome ejection remains to be identified.

### Role of newly synthesized histones in parental histone recycling

To maintain correct nucleosome density on the replicated daughter DNA strands, nucleosomes are assembled from the pool of recycled parental histones and newly synthesized histones. Assuming that all parental histones are reinstated during replication, an equal amount of newly synthesized histones needs to be delivered into the nucleus to restore chromatin structure. This high demand for canonical core histones during S phase is fulfilled through rapid expression of multicopy histone genes, induced at the onset of replication and tightly regulated throughout the cell cycle (Marzluff et al., 2008). Because histones are highly basic proteins, and so have the potential to bind non-specifically to negatively charged macromolecules, such as DNA and RNA, they are escorted throughout their cellular life by dedicated networks of chaperone proteins (Gurard-Levin et al., 2014). Histone chaperones ensure their correct folding, control their traffic within the cell (such as nuclear import, nucleosome assembly and histone degradation) and assist nucleosome dynamics. The deficit or excess of canonical histones was found to inhibit DNA replication and lead to genomic instability in yeast and mammalian cells (Groth et al., 2007; Gunjan and Verreault, 2003; Han et al., 1987; Kim et al., 1988; Mejlvang et al., 2014; Nelson et al., 2002).

*Xenopus* eggs naturally contain high amounts of histones because they need to support the first twelve rounds of DNA replication prior to the midblastula transition, when transcription is initiated (Newport and Kirschner, 1982). Consequently, egg extracts have a significantly higher concentration of ‘free’ histones than an equivalent extract of somatic cells. The quantitative analysis of the replication fork collision with nucleosomes in these extracts revealed that nucleosome/histone eviction is the dominant outcome, approximately three times more likely than parental histone transfer (Figure 6). Interestingly, extracts depleted of a large proportion of newly synthesized histones promoted efficient parental histone recycling, increasing its likelihood to ∼3:1 over histone eviction. Supplementation of depleted extracts with recombinant histones reversed this effect, resulting in nucleosome/histone eviction prevalence over localized histone transfer, at ∼2:1 likelihood ratio.

Our analysis clearly demonstrates that the efficiency of localized parental histone recycling depends on the concentration of newly synthesized histones. We interpret these results with the following molecular model (Figure 7). At low concentrations of free histones, the majority of parental histones is locally recycled. Upon nucleosome disassembly ahead of the replication fork, parental histones are released from the DNA and remain in the vicinity of the replisome, through a concerted action of histone chaperones and replisome components, which finally deposit parental histones on the daughter DNA. When the concentration of newly synthesized histones is high, most parental histones are released into the milieu and do not get incorporated into replicated DNA. The most probable explanation for such behavior is that the free histones exchange with their parental counterparts *en route* from parental to nascent DNA. Although less likely, we cannot rule out the possibility that the pathway of newly synthesized histone deposition takes over in conditions of excessive histone provision and inhibits localized parental histone recycling.

**Figure 7.**
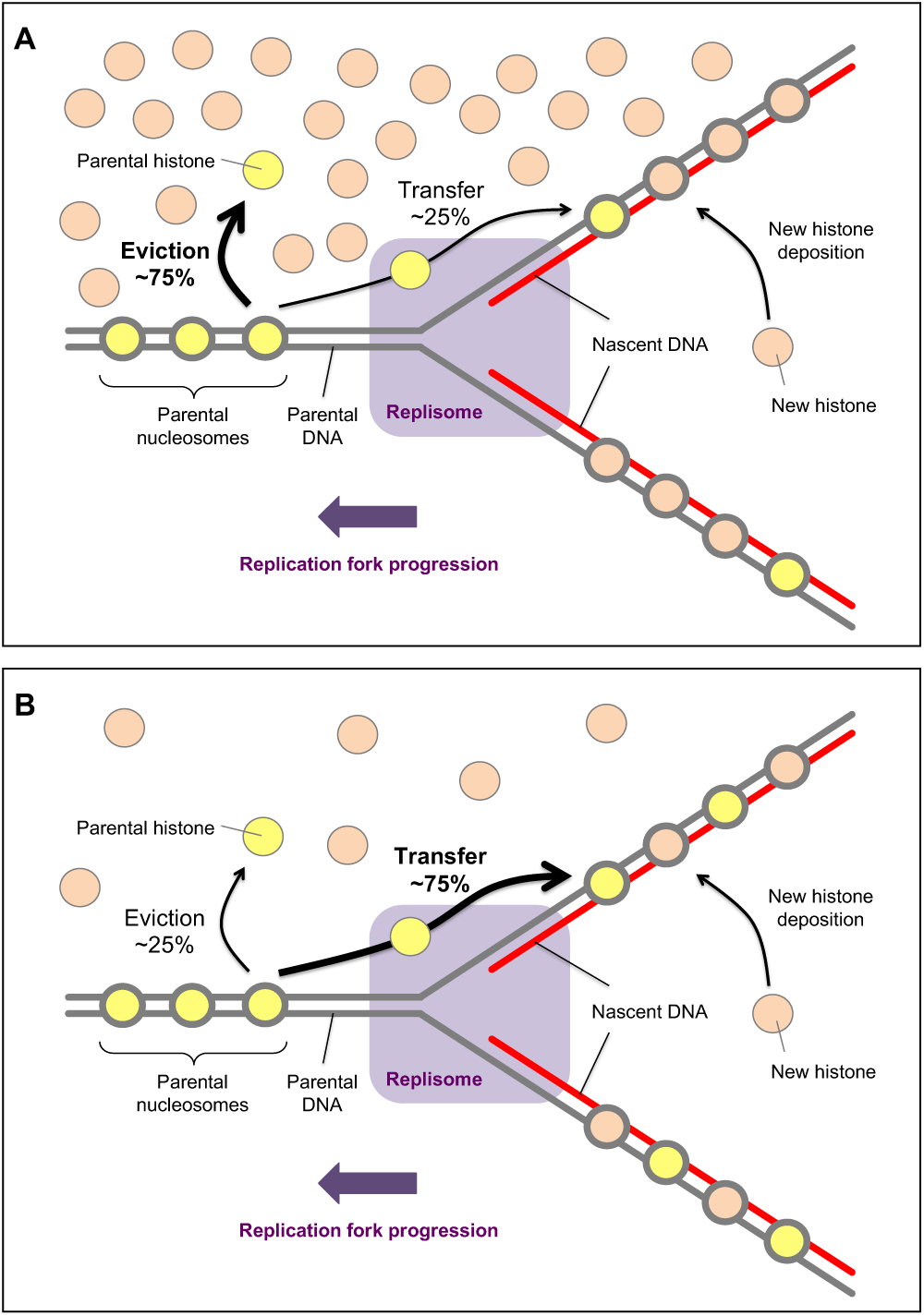
Model of parental histone transfer at high and low concentrations of newly synthesized histones. (A) At high concentrations of free histones, upon the encounter with the replication machinery, the majority of parental histones are evicted from the DNA and released into the histone pool. (B) When the concentration of newly synthesized histones is low, the majority of parental histones is recycled at the replication fork. Upon nucleosome disassembly ahead of the replication fork, parental histones are released from the DNA but are kept in the vicinity of the replisome, most likely through concerted action of histone chaperones and replisome components. Parental histones are rapidly ushered behind the replication fork where they are deposited onto daughter strands.

### Consequences for epigenetic inheritance

Epigenetic changes are heritable alterations to gene expression profiles occurring without modifications to the primary structure of DNA. Chromatin is divided into domains that either facilitate transcription (euchromatin or open chromatin) or repress it (heterochromatin or compacted/closed chromatin). Nucleosomes in eu- and heterochromatin carry specific histone PTMs, which modulate the structure and dynamics of these chromatin structures (Stillman, 2018). Thus, in order to maintain the transcriptional program of the cell, chromatin structures and their associated histone PTMs, must be faithfully transmitted to daughter cells, through a process generally referred to as epigenetic inheritance. The key question in the field of epigenetics is whether localized parental histone recycling at the replication fork drives the transgenerational transmission of PTMs.

Through the use of ChOR-seq (chromatin occupancy after replication) in HeLa cells, it was recently reported that accurate parental histone redeposition preserves positional information and allows PTM transmission to daughter cells (Reveron-Gomez et al., 2018). This behavior was observed for histone modifications marking transcriptionally active chromatin (tri-methylation of histone H3 at lysine 4, lysine 36 and lysine 79; H3-K4me3, H3-K36me3 and H3-K79me3, respectively) as well as transcriptionally silent chromatin (tri-methylation of histone H3 at lysine 27, H3-K27me3). Interestingly, microscopic tracking of parental histone H3 variants through cell division in HeLa cells (Clement et al., 2018) demonstrated that H3.3, marking early-replicating chromatin (known to be transcriptionally active), was lost at a faster rate than H3.1, which is associated with late-replicating chromatin (characteristic of transcriptionally repressed domains) (Rhind and Gilbert, 2013). Furthermore, a locus-specific, proximity-dependent histone labelling-based study in mouse embryonic stem cells has revealed that only the repressed chromatin domains are preserved through local recycling of parental histones (Escobar et al., 2019). In the case of active chromatin, the local redeposition of parental histones was absent in this study, leading the authors to suggest that the associated histone PTMs are not epigenetic and function solely to facilitate transcription. Indeed, the concept that PTMs can be epigenetically inherited through localized histone recycling originated from studies of tri-methylation of histone H3 on lysine 9 (H3-K9me3) and lysine 27 (H3-K27me3), which are associated with constitutive and facultative heterochromatin, respectively (Reinberg and Vales, 2018). The model suggests that these PTMs segregate to daughter strands during DNA replication, producing a landscape where nucleosomes containing parental histones with the PTMs are adjacent to naïve nucleosomes, composed of newly synthesized histones. The protein machineries responsible for depositing H3-K9me3 and H3-K27me3 operate by using a specific ‘read and write’ mechanism, whereby they first bind to the existing PTM on a recycled parental histone, which in turn stimulates the modification of neighboring naïve nucleosomes. The ‘read and write’ mechanism ensures that repressive chromatin domains are efficiently reestablished in daughter cells.

Therefore, a major question in the field is what molecular mechanism could lead to different patterns of parental histone recycling at the replication fork for eu- and heterochromatin. A possible explanation is that there are PTM-specific chaperones directing parental histones either for local transfer onto daughter strands (in the case of PTMs associated with heterochromatin) or into the pool of soluble histones (in the case of PTMs associated with euchromatin). This model however seems unlikely given that histones without any posttranslational modifications were found to efficiently transfer onto daughter strands in a replication system reconstituted from yeast purified proteins (in the absence of soluble histones) (Kurat et al., 2017). Another reason for the observed discrepancy in local histone conservation between transcriptionally active and repressed chromatin could be the rate of replication, known to be faster for early-replicating euchromatin than late-replicating heterochromatin (Rhind and Gilbert, 2013). Our analysis of the replication fork collision with nucleosomes shows that the velocity of the progressing replication fork has no influence on the collision outcome (Figure 6G), rendering this explanation less probable. Based on our findings that the efficiency of parental histone transfer depends on the concentration of newly synthesized histones (Figures 6 and 7), we propose an alternative molecular mechanism for the selective epigenetic conservation across chromatin domains in which differential levels of accessible free histones are used to prevent local histone recycling in euchromatin but promote it in heterochromatic regions. Rapid histone biosynthesis is activated at the beginning of S phase and persists at high levels until the end of S phase, when DNA replication is halted (Marzluff et al., 2008). Transcriptionally active and silenced chromatin domains display distinct spatial segregation in the nucleus (Bonev and Cavalli, 2016; Solovei et al., 2016; van Steensel and Belmont, 2017) and their replication is separated in time. Recent studies show that associations between heterochromatic regions, most likely driven by HP1*α* (Larson et al., 2017; Strom et al., 2017), lead to phase separation of active and repressed chromatin, whereas euchromatic interactions are dispensable for compartmentalization (Falk et al., 2019). We speculate that the phase boundary could act as a selective barrier to histones and/or the associated chaperones, and thus provide distinct regions of histone accessibility within the nucleus during replication. Phase separated heterochromatin domains would replicate under conditions of limited provision of newly synthesized histones, ensuring efficient localized parental histone transfer at the replication fork, and so its epigenetic inheritance. In the case of transcriptionally active euchromatin, the high local concentration of newly synthesized histones would lead to dispersed redistribution of parental and newly synthesized histones on daughter strands.

## ACKNOWLEDGEMENTS

We thank Daniel Maskell for H3-K36C histone octamer. We thank Johannes Walter for pET28-Fen1-KikGR plasmid, and Gheorghe Chistol and Livio Dukaj for helpful discussions on Fen1-KikGR experiments. We thank Hironori Funabiki and Christian Zierhut for advice on histone depletion from *Xenopus* egg extracts. We thank Geneviève Almouzni for anti-Asf1 antibody. This work was supported by the Francis Crick Institute, which receives its core funding from Cancer Research UK (FC001221), the UK Medical Research Council (FC001221), and the Wellcome Trust (FC001221). H.K. was supported by JSPS (JP17H01417, and JP18H05527) and JST-CREST (JPMJCR16G1).

## AUTHOR CONTRIBUTIONS

D.T.G and H.Y. designed the experiments and prepared the manuscript. D.T.G. performed and analyzed the experiments. S.X. purified p27^Kip^ and geminin. H.K. supplied anti-H4-K12Ac antibody.

## VIDEO TITLES AND LEGENDS

**Video 1. Related to Figure 2. Singly-tethered low-density λ nucleosomes containing H3-K36C^Cy5^ (yellow) imaged in buffer under ‘no-flow’ conditions followed by 50 μl/min flow**. Without buffer flow low-density nucleosomes on a single *λ* DNA molecule appear as a diffraction-limited spot of fluorescence. The molecule unfolds and stretches under flow, unveiling nucleosomes distributed along *λ* as ‘beads-on-a-string’.

**Video 2. Related to Figure 2. Doubly-tethered high-density λ nucleosomes containing H3-K36C^Cy5^ (yellow) imaged in buffer under ‘no-flow’ conditions followed by 50 μl/min flow**. Doubly-tethered *λ* molecule saturated with nucleosomes does not stretch under flow, and thus appears as a diffraction-limited spot of fluorescence throughout the movie.

Video 3. Related to Figure 3B. Stretched λ nucleosomes containing H2A-K119C^Cy5^ (yellow) during incubation in HSS (3-14min).

Video 4. Related to Figure 3B. Stretched λ nucleosomes containing H2B-T112C^A647^ (yellow) during incubation in HSS (3-14min).

Video 5. Related to Figure 3C. Stretched λ nucleosomes containing H3-K36C^Cy5^ (yellow) during incubation in HSS (3-14min).

Video 6. Related to Figure 3C. Stretched λ nucleosomes containing H3-T80C^A647^ (yellow) during incubation in HSS (3-14min).

Video 7. Related to Figure 3C. Stretched λ nucleosomes containing H4-E63C^A647^ (yellow) during incubation in HSS (3-14min).

**Video 8. Related to Figure 4A. Example of nucleosome-fork collision resulting in nucleosome (histone) eviction**. Nucleosome/histone eviction is manifested by the loss of histone fluorescence (H4-E63C^A647^; yellow) at the point of collision with the progressing replication fork (Fen1-KikGR; red).

**Video 9. Related to Figure 4B. Example of nucleosome-fork collision resulting in histone transfer behind the replication fork**. Histone transfer is observed when the histone-associated fluorescence H4-E63C^A647^; yellow) is retained and incorporated into the track of replicated DNA (Fen1-KikGR; red).

**Video 10. Related to Figure 4C. Example of nucleosome-fork collision resulting in nucleosome/histone sliding**. Nucleosome (histone) sliding is observed when the histone-associated fluorescence (H4-E63C^A647^; yellow) moves together with the tip of the replication bubble (Fen1-KikGR; red).

**Video 11. Related to Figure 4D. Example of nucleosome-fork collision resulting in replication fork stalling**. Replication fork stalling occurs when nucleosome constitutes a roadblock preventing the replication fork from further movement and is manifested by an arrested tip of the replication bubble (Fen1-KikGR; red) next to a static histone signal (H4-E63C^A647^; yellow).

**Video 12. Related to Figure 5A. Example of nucleosome-fork collision resulting in nucleosome/histone sliding followed by eviction**. H4-E63C^A647^ histones are shown in yellow and the Fen1-KikGR-decorated replication bubble is shown in red.

**Video 13. Related to Figure 5B. Example of nucleosome-fork collision resulting in replication fork stalling followed by nucleosome/histone eviction**. H4-E63C^A647^ histones are shown in yellow and the Fen1-KikGR-decorated replication bubble is shown in red.

**Video 14. Related to Figure 5C. Example of nucleosome-fork collision resulting in nucleosome/histone sliding followed by histone transfer**. H4-E63C^A647^ histones are shown in yellow and the Fen1-KikGR-decorated replication bubble is shown in red.

**Video 15. Related to Figure 5D. Example of nucleosome-fork collision resulting in replication fork stalling followed by histone transfer**. H4-E63C^A647^ histones are shown in yellow and the Fen1-KikGR-decorated replication bubble is shown in red.

**Video 16. Related to Figure 5E. Example of nucleosome-fork collision resulting in nucleosome/histone sliding followed by replication fork stalling**. H4-E63C^A647^ histones are shown in yellow and the Fen1-KikGR-decorated replication bubble is shown in red.

**Video 17. Related to Figure 5F. Example of nucleosome-fork collision resulting in replication fork stalling followed by nucleosome/histone sliding**. H4-E63C^A647^ histones are shown in yellow and the Fen1-KikGR-decorated replication bubble is shown in red.

## METHODS

### Preparation of biotinylated λ DNA

Singly-biotinylated *λ* DNA was prepared as described in (Yardimci et al., 2012). Doubly-biotinylated *λ* DNA was prepared by mixing 80 μM biotin-14-dCTP (Invitrogen; 19524-016), 80 μM biotin-14-dATP (Invitrogen; 19518-018), 100 μM dTTP (Thermo Fisher Scientific; R0171), 100 μM dGTP (Thermo Fisher Scientific; R0161), 130 ng/μl *λ* DNA (NEB; N3011) and 0.05 U/μl of Klenow fragment (NEB; M0212S) in provided Klenow buffer. Mixture was incubated at 37°C for 30 minutes, followed by 15 minutes at 70°C. *λ* DNA was purified using QIAquick PCR Purification Kit (QIAGEN; 28104) and stored at 4°C. This method introduces multiple biotins at each end of *λ* DNA (7 biotins at the left end and 4 biotins at the right end, assuming 100% incorporation of biotinylated dNTPs).

### Histone labelling under denaturing conditions

Purified, recombinant *Xenopus* histones were purchased from The Histone Source, Protein Expression and Purification Facility, Colorado State University, and their correct molecular mass was verified by mass spectrometry (Proteomics Science Technology Platform, Francis Crick Institute). Histones H2A-K119C, H2B-T112C, H3-C110A-T80C and H4-E63C were labelled with either Cy®5 maleimide (GE Heathcare; PA25031) or Alexa Fluor^TM^ 647 C_2_ maleimide (Thermo Fisher Scientific; A20347) using thiol-modification of engineered cysteines. Prior to labelling, histones were reduced and denatured in 20 mM Tris-HCl pH 7.5 (Sigma; T1503; and Fisher Scientific; 10316380), 10 mM Tris(2-carboxyethyl)phosphine (TCEP; Sigma; 646547) and 7M guanidine hydrochloride (Sigma; 50940) for 30 minutes at room temperature. Each denaturing reaction contained a chosen histone at a concentration of 150 μM in a total volume of 250 μl (equivalent to approximately 0.5 mg of histone). One vial of Cy®5 maleimide or 0.5 mg of Alexa Fluor^TM^ 647 C_2_ maleimide was dissolved in 50 μl of anhydrous dimethyl sulfoxide (DMSO; Invitrogen; D12345) and then mixed dropwise with 250 μl of the denatured histone solution. Labelling reactions were carried out for 2.5–3 hours at room temperature and protected from light. β-mercaptoethanol (Sigma; 101458612) was added to a labelling reaction at a 100-fold molar excess of the dye to consume any unreacted species. The quenched reaction was used immediately to refold histone octamer.

### Histone octamer refolding and purification

Histone octamer refolding protocol was adapted from (Dyer et al., 2004). Histones were individually reduced and denatured in 20 mM Tris-HCl pH 7.5, 10 mM β-mercaptoethanol and 7M guanidine hydrochloride for 3 hours at room temperature. Each denaturing reaction contained a chosen histone at a concentration of 150 μM in a total volume of 250 μl (equivalent to approximately 0.5 mg of histone). Denatured histones H2A, H2B, H3 and H4, were mixed at equimolar ratios and adjusted to a total protein concentration of 1 mg/ml with unfolding buffer (20 mM Tris-HCl pH 7.5, 10 mM β-mercaptoethanol and 7M guanidine hydrochloride). For labelled octamer refolding, a quenched labelling reaction was used instead of a wild-type denatured histone. Denatured histone mix was loaded into a MaxiGeBaFlex dialysis tube (Generon; D045; 8 kDa molecular weight cutoff; 2–3 ml capacity) and dialyzed at 4°C against 2 l of 10 mM Tris-HCl pH 7.5, 1 mM ethylenediaminetetraacetic acid disodium salt dihydrate (EDTA; Sigma; E5134), 5 mM β-mercaptoethanol and 2 M NaCl (Sigma; S9888). Refolding buffer was changed at least three times for unlabelled octamer and four times for fluorescently labelled octamers (1^st^ – overnight, 2^nd^ – 8 hours, 3^rd^ – overnight, 4^th^ – 8hrs).

Refolded histone mixture was recovered from the dialysis device and concentrated to approximately 0.3 ml using VivaSpin 500 centrifugal concentrator (Sartorius; VS0121; 30 kDa molecular weight cutoff; PES) at 2°C, 15,000 x g. Concentrated sample was resolved on a Superdex 200 Increase GL10/300 column (GE Heathcare; 28-9909-44), over 1.1 column volume of refolding buffer (10 mM Tris-HCl pH 7.5, 1 mM EDTA, 5 mM β-mercaptoethanol and 2 M NaCl) at 0.3 ml/min flow rate, 4°C. Fractions containing stoichiometric octamer, as verified by SDS-PAGE, were pooled and concentrated using VivaSpin 500 centrifugal concentrator (30 kDa molecular weight cutoff; PES). Octamer concentration and labelling efficiency were estimated spectrophotometrically from the absorbance measurement at 276 and 650 nm. Octamer was flash-frozen in liquid nitrogen and stored at -80°C.

### Histone octamer labelling under native conditions

Histone octamer containing H3-K36C^Cy5^ was prepared by thiol-modification under native conditions. Octamer containing unlabelled H3-K36C was refolded and purified as described above, but the unfolding and refolding buffers contained TCEP, instead of β-mercaptoethanol, as a reducing agent. 0.5 mg of octamer was adjusted to a concentration of 1 mg/ml with refolding buffer. One vial of Cy®5 maleimide was dissolved in 50 μl of anhydrous DMSO and then mixed dropwise with the octamer solution. Labelling reactions were carried out overnight at 2°C, protected from light. β-mercaptoethanol (Sigma; 101458612) was added to a labelling reaction at a 100-fold molar excess of the dye to quench any unreacted species. Excess dye was removed using Micro Bio-Spin P-30 Columns (Bio-Rad; 7326202), pre-equilibrated with refolding buffer. Octamer concentration and labelling efficiency were estimated spectrophotometrically from the absorbance measurement at 276 and 650 nm. Octamer was flash-frozen in liquid nitrogen and stored at -80°C.

### Nucleosome reconstitution

Nucleosome reconstitution was performed by NaCl gradient dialysis method. For each reconstitution reaction, 1 μg of DNA was mixed with a desired molar excess of histone octamer (from 0 to 300 for *λ* DNA) in 10 mM Tris-HCl pH 7.5, 1 mM EDTA and 2 M NaCl, to a final volume of 100 μl, and incubated on ice for 30 min. Samples were then transferred into a Slide-A-Lyzer® MINI dialysis units (Thermo Scientific; 96570) and dialyzed overnight against 1 l of 10 mM Tris-HCl pH 7.5, 1 mM EDTA and 1 M NaCl. Second dialysis was performed for 8 hours against 1 l of 10 mM Tris-HCl pH 7.5, 1 mM EDTA and 0.75 M NaCl, before the final overnight dialysis against 1 l of 10 mM Tris-HCl pH 7.5, 1 mM EDTA and 20 mM NaCl. Reconstituted nucleosomes were recovered from the dialysis devices and stored at 4°C. Samples containing fluorescently labelled histones were protected from light at each step.

### Electrophoretic mobility shift assay (EMSA)

100 ng of naked *λ* DNA or *λ* nucleosomes in 10 mM Tris-HCl pH 7.5, 1 mM EDTA, 20 mM NaCl, 10 % glycerol (Fisher Scientific; BP229-1) were resolved on a 0.5 % agarose (Denville Scientific Inc.; CA3510-8) gel in 20 mM Tris and 20 mM boric acid (Fisher Chemical; B/3800/53) for 120 minutes at 100 V. After electrophoresis, DNA was stained with SYBR^TM^ Gold nucleic acid gel stain (Thermo Fisher Scientific; S11494), following the manufacturer’s protocol. Gels were imaged using fluorescent image analyzer FLA-5000 (Fujifilm). Samples containing fluorescently labelled histones were protected from light at each step.

### Native micrococcal nuclease (MNase) protection assay

100 ng of naked *λ* or *λ* nucleosomes in 10 mM Tris-HCl pH 7.5, 1 mM EDTA and 20 mM NaCl were supplemented with micrococcal nuclease buffer (NEB; M0247S), following the manufacturer’s instructions, and then digested with 10 GU of micrococcal nuclease (MNase; NEB; M0247S) for 10 minutes at room temperature. Digest was quenched by adding EDTA to a concentration of 25 mM and 10 % glycerol was used as a loading agent. Digested samples were resolved on a 1.5 % agarose gel in 20 mM Tris and 20 mM boric acid for 120 minutes at 100 V. After electrophoresis, DNA was stained with SYBR^TM^ Gold nucleic acid gel stain and imaged using fluorescent image analyzer FLA-5000. Samples containing fluorescently labelled histones were protected from light at each step.

### Denaturing micrococcal nuclease (MNase) protection assay

In the denaturing MNase protection assay, samples were prepared, digested and quenched as described for native assay. Upon quenching with EDTA, each sample was supplemented with sodium dodecyl sulfate (SDS; Sigma; 436143) to a concentration of 0.8 % and 0.8 U of proteinase K (NEB; P8107S). Protein digest was conducted at 37°C for 1 hour. Samples were supplemented with glycerol to 10 % and resolved on a 1.5 % agarose gel in 100 mM Tris, 100 mM boric acid and 2 mM EDTA (TBE). DNA was stained with SYBR^TM^ Gold nucleic acid gel stain and imaged using fluorescent image analyzer FLA-5000.

### *Xenopus laevis* egg extracts preparation

High speed supernatant (HSS) and nucleoplasmic extract (NPE) were prepared as described previously (Lebofsky et al., 2009) and stored at -80°C. Prior to both bulk and single-molecule replication assays, each 33 μl aliquot of HSS was supplemented with 250 ng of nocodazole (Sigma; M1404) and 1 μl of an ATP regeneration system, containing 650 mM phosphocreatine (Sigma; P7936), 65 mM ATP (pH 7.0; Sigma; A2754) and 0.161 mg/ml creatine phosphokinase (Sigma; C3755). Similarly, each 16 μl aliquot of NPE was supplemented with 0.5 μl of ATP mix. Activated extracts were centrifuged for 5 minutes at 16,000 x g, room temperature, and used in replication assays.

### Histone depletion from *Xenopus* egg extracts

50 μl (bed volume) of protein A sepharose (PAS; GE Healthcare; GE17-1279-01), pre-washed with ice-cold phosphate-buffered saline (PBS; Gibco; 70011044; six times with 300 μl), was mixed with 300 μl of a 1.6 mg/ml anti-H4-K12Ac antibody solution in PBS, and then incubated overnight at 4°C, 20 rpm. PAS loaded with an antibody was washed four times with 300 μl of cold PBS and three times with 300 μl of cold ELB by centrifugation. 200 μl of HSS-NPE mix (extracts were not supplemented with ATP mix but nocodazole was added into HSS to prevent microtubule polymerization) at 1:1 volume ratio was next mixed with 50 μl (bed volume) of antibody-loaded PAS and incubated for 1 hour at 4°C, 20 rpm. Extracts were separated from PAS by spinning through a nitex column, as described in (Lebofsky et al., 2009). Cleared extracts were mixed with 34 μl (bed volume) of PAS, pre-washed with cold PBS (six times with 300 μl) and ELB (three times with 300 μl), and incubated for 45 minutes at 4°C, 20 rpm. This step ensures that any leftover antibody is captured and removed from extracts. Finally, depleted extracts were clarified on a nitex spin column and either used immediately in replication assays or snap-frozen in liquid nitrogen and stored at -80°C.

### Bulk replication assay

Naked pBRII (pBlueScript II; Agilent Technologies; 212205) plasmid and pBRII containing fluorescent nucleosomes (at a saturation level equivalent with *λ* nucleosomes in single molecule replication assays) labelled at H3-K36C^Cy5^ or H4-E63CA^647^ were adjusted to a DNA concentration of 18 ng/μl with egg lysis buffer (ELB; 2.5 mM MgCl_2_, 50 mM KCl, 10 mM HEPES-KOH, pH 7.7; Sigma M8266; Sigma P9333; Sigma H3375), supplemented with ATP mix (1 μl per 16 μl of DNA in ELB), and then mixed at 1:1 volume ratio with activated HSS. Equivalent reactions were set up with HSS supplemented and preincubated (5 minutes at room temperature) with 4 μM geminin, as replication-negative controls. All samples were incubated for 15 minutes at room temperature to promote origin licensing. 16 μl of activated NPE was supplemented with 0.2 μl of 10 mCi/ml [*α*-^32^P]dATP (3 kCi/mmol; Perkin Elmer; BLU512H250UC). [*α*-^32^P]dATP gets incorporated into nascent DNA strands during replication, and thus allows to track the progress of replication in time. At 8, 15 and 30 minutes after NPE was introduced, a 2.5 μl aliquot of a replication reaction was stopped by mixing in 5.0 μl of solution containing 25 mM Tris-HCl pH 8.0, 2% SDS, 75 mM EDTA and 8 U/ml proteinase K, and incubated at 37°C for 1 hour. Replication reactions were separated on a 0.8 % agarose gel in TBE at 90 V, room temperature. Gel was dried and visualized using fluorescent image analyzer FLA-5000 in a phosphorescence mode.

### Expression and purification of Fen1-KikGR

Fen1-KikGR was expressed and purified from *E. coli* with some modifications to the protocol described in (Loveland et al., 2012). BL21-CodonPlus (DE3)-RIPL competent cells (Agilent Technologies; 230280) were transformed with the pET28-Fen1-KikGR expression plasmid and selected on LB-agar plates supplemented with 50 μg/ml of kanamycin (Sigma; BP861). A single colony was used to inoculate 50 ml of selective LB medium. After overnight incubation at 37°C on a shaker (220 rm), 5 ml of this starter culture was used to inoculate 500 ml of fresh selective LB medium. Cultures were grown at 37°C, with shaking, until the optical density at 600 nm reached 0.6 and were then cooled down to 20°C in a water bath. Isopropyl-β-D-thiogalactoside (IPTG; Fisher Scientific; 10725471) was added to a final concentration of 0.5 mM and cultures were grown overnight at 20°C and protected from light. Cells were harvested by centrifugation for 15 minutes at 5’000 x g, 4°C.

Cell pellets from 2 l of culture were resuspended in 20 mM Tris-HCl pH 8.0, 1 M NaCl, 1 mM 1,4-dithiothreitol (DTT; Sigma; 10197777001), 50 mM imidazole (Sigma; I5513) to a final volume of 40 ml and lysed by sonication (60 pulses of 3 seconds at 70 W with 7 second pauses between pulses). The crude extract was cleared from insoluble cell debris by centrifugation for 45 minutes at 45,000 x g at 4°C. The supernatant was filtered through a 0.22 μm PES membrane (Merck Millipore; SLGP033RS), supplemented with EDTA-free protein inhibitor cocktail (Roche; 5056489001) and loaded onto a 5 ml HisTrap HP column (GE Healthcare; 17-5248), pre-equilibrated with 20 mM Tris-HCl pH 8.0, 1 M NaCl, 1 mM DTT, 50 mM imidazole. Proteins were eluted with a linear gradient of imidazole from 50 to 500 mM over 10 column volumes at 4°C. Eluted fractions were analyzed by SDS-PAGE and those containing Fen1-KikGR were pooled and dialyzed overnight at 4°C against 10 mM HEPES-KOH pH 7.7, 2.5 mM MgCl_2_, 300 mM NaCl, 1 mM EDTA, 1 mM DTT, 10 % glycerol. Dialyzed protein solution was concentrated using VivaSpin 20 centrifugal concentrators, 10 kDa molecular weight cutoff (Sartorius; VS2002). The final concentration of purified Fen1-KikGR was estimated spectrophotometrically from the absorbance measurement at 280 and 505 nm. 3 μl aliquots of Fen1-KikGR were flash-frozen in liquid nitrogen and stored at -80°C.

### Single-molecule replication assay

Microfluidic flow cells with PEGylated and streptavidin-functionalized glass surface were prepared as described previously (Yardimci et al., 2012). Flow cells were mounted on a Nikon Eclipse Ti motorized inverted microscope, equipped with a 100x high numerical aperture TIRF objective (SR Apo TIRF 100x 1.49 Oil; Nikon), the Perfect Focus System and supported by LU-N4 laser unit (Nikon), providing four lasers: 405, 488, 561 and 640 nm (15 mW output power at the fiber end). Images were recorded using a 512 x 512 pixel, back illuminated, electron-multiplying charge-coupled-device camera (iXon DU-987, Andor Technology; 3 MHz pixel readout rate, 14 bit digitization and 300x electron multiplier gain) controlled by NIS-Elements software (Nikon). The pixel size was 160 x 160 nm. All buffers and solutions were thoroughly degassed immediately before use. Flow was controlled by an automated syringe pump (Pump 11 Elite; Harvard Apparatus; 70-4505). All experiments were conducted at room temperature.

Prior to DNA immobilization, microfluidic channels were washed with blocking buffer containing 20 mM Tris pH 7.5, 50 mM NaCl, 2 mM EDTA and 0.2 mg/ml BSA (albumin from bovine serum; Sigma; A7906). For immobilization of singly-biotinylated *λ* nucleosomes, 125 μl of DNA or nucleosome solution at a concentration of 0.1 ng/μl in blocking buffer were passed through the channel at a flow rate of 25 μl/min. DNA was incubated in the channel for 10 minutes and any unbound molecules were removed by washing with 250 μl of blocking buffer at 50 μl/min flow rate. Doubly-biotinylated naked *λ* or *λ* nucleosomes were immobilized by passing through 500 μl of DNA or nucleosome solution at a concentration of 0.1 ng/μl in blocking buffer at a flow rate of 100 μl/min. This procedure immobilizes *λ* DNA and *λ* nucleosomes to approximately 70 % of their respective, maximally stretched contour lengths. Cy5 or Alexa Fluor 647 labelled histones within immobilized nucleosomes were imaged using 640-nm laser at 10 % power, 100 ms exposure time and ZT405/488/561/647rpc dichroic (Chroma). Tethered DNA molecules were stained with 5 nM SYTOX^TM^ Orange (Thermo Fisher Scientific; S11368) in blocking buffer and imaged using 560-nm laser at 5 % power, 100 ms exposure time and ZT405/488/561/647rpc dichroic. To remove SYTOX Orange, flow cell was washed extensively with blocking buffer, 0.5 – 1.0 ml at a flow rate of 50 μl/min. Immediately before licensing, ELB supplemented with casein (Sigma; C4765) and BSA to a final concentration of 1 mg/ml was introduced into the channel at 25 μl/min for 3 minutes.

For licensing of the immobilized DNA, an aliquot of activated and span down HSS (see *Xenopus laevis* egg extracts preparation) was transferred to a fresh tube, supplemented with a short linear ‘carrier’ DNA (pre-annealed oligos 5’-GCA GCA ACA GAA GCC ATG GAT GCC CTG AC-3’ and 5’-GTC AGG GCA TCC ATG GCT TCT GTT GCT GC-3’) to a concentration of 10 ng/ul and incubated for 5 minutes. HSS was introduced into the channel at a flow rate of 10 ul/min over 2.5 minutes and incubated for further 12.5 minutes. During licensing in HSS, Cy5 or Alexa Fluor 647 labelled histones were imaged using 640 nm laser at 5 % power, 100 ms exposure time and ZT405/488/561/647rpc dichroic (Chroma). Images were collected for 25 different fields of view (5 x 5 grid; 512 x 512 pixel per field of view) at a 11-seconds interval between frames.

While the licensing reaction was taking place, replication extracts were prepared by mixing activated HSS, NPE and ELB at 1:1:1 volume ratio. The replication mix was further supplemented with the pBRII plasmid to a final concentration of 5 ng/μl, Fen1-KikGR to 2.5 μM and oxygen scavenging system (i.e. glucose to 40 mM, pyranose oxidase to 2.5 U/ml and catalase to 120 U/ml; Sigma G8270; Sigma P4234-250UN; Sigma C30-100MG; (Swoboda et al., 2012)). For unrestricted origin firing (replication from multiple origins), 40 μl of this mix was drawn into the channel at a 10 μl/min flow rate. To achieve replication from single origins, the mix was split into two 20 μl aliquots. One aliquot was immediately drawn into the channel at 10 μl/min for 2 minutes to initiate replication of licensed and immobilized DNA molecules. The other aliquot was supplemented with p27^Kip^, a Cdk2 inhibitor, to a concentration of 0.1 μg/μl and introduced into the channel when about one or two origins per template fired; typically, between 4 to 8 minutes from the moment the first extract was drawn in. During replication, Cy5 or Alexa Fluor 647 labelled histones were imaged using 640-nm laser at 5 % power, 100-ms exposure time and ZT405/488/561/647rpc dichroic. Fen1-KikGR-decorated replication bubbles were imaged using 488-nm laser at 5 % power, 100-ms exposure time and ZT405/488/561/647rpc dichroic. Unless stated otherwise, images were collected for 36 different fields of view (6 x 6 grid; 512 x 512 pixel per field of view) at a 1-minute interval between frames.

For replication in extracts depleted of endogenous histones H4 and H3, DNA template immobilization and licensing were conducted as described above. 16 μl of H4(H3)-depleted HSS-NPE mix was supplemented with 0.5 μl of ATP mix and centrifuged for 5 minutes at 16,000 x g, room temperature. The activated extract mix was next transferred to a fresh tube and supplemented with pBRII to a final concentration of 5 ng/μl, Fen1-KikGR to 2.5 μM and oxygen scavenging system (i.e. glucose to 40 mM, pyranose oxidase to 3 U/ml and catalase to 90 U/ml). In the case of replication experiments in extracts depleted of endogenous histones but supplemented with recombinant histones, the activated mix was additionally supplemented with histones H3 and H4 to a final concentration of 20 μM. The volume was adjusted to 20 μl with ELB, the mixture was drawn into the channel and imaging was conducted as described for undepleted extracts. Depleted extracts showed lower overall origin firing efficiency, relative to undepleted extracts, and so did not require p27^Kip^ supplementation for individual bubble growth tracking during replication.

For replication in the absence of Fen1-KikGR, DNA template immobilization and licensing were conducted as described above. Replication extracts were prepared by mixing activated HSS, NPE and ELB at 1:1:1 volume ratio. The replication mix was further supplemented with pBRII to a final concentration of 5 ng/μl, digoxygenin-11-dUTP (dig-dUTP; Roche; 11093088910) to 1.7 mM and oxygen scavenging system (i.e. glucose to 40 mM, pyranose oxidase to 3 U/ml and catalase to 90 U/ml). The mix was split into two 20 μl aliquots. One aliquot was immediately drawn into the channel at 10 μl/min for 2 minutes to initiate replication. The other aliquot was supplemented with p27^Kip^ to a concentration of 0.1 μg/μl and introduced into the channel at 9 minutes from the moment the first extract was drawn in. Replication elongation was allowed to proceed for next 31 minutes before a buffer containing 20 mM Tris pH 7.5, 10 mM EDTA and 0.5 M NaCl was flown in at a rate of 20 μl/min over 10 minutes to wash out the extracts. The flow cell was next washed with 250 μl of blocking buffer at 50 μl/min flow rate. 350 μl of a 0.2 ng/μl solution of fluorescein labelled anti-digoxigenin Fab fragments from sheep (anti-dig Ab^Fluor^; Roche; 11207741910) in blocking buffer, supplemented with 1 mg/ml of casein and 1 mg/ml of BSA, was introduced into the chamber at a flow rate of 10 μl/min. The flow cell was next washed with 100 μl of blocking buffer at 20 μl/min flow rate. Finally, *λ* DNA was stained with a 5 nM SYTOX Orange solution in blocking buffer, drawn into the cell at a rate of 20 μl/min. Cy5-labelled histones were imaged using 640-nm laser at 5 % power, 100-ms exposure time and ET700/50m emission filter (Chroma). Nascent DNA, decorated with anti-dig Ab^Fluor^, was imaged using 488-nm laser at 2 % power, 100-ms exposure time and ET525/50m emission filter (Chroma). Sytox was excited using 561-nm laser at 5 % power, 100-ms exposure time and ET600/50 m emission filter (Chroma).

### Single-molecule data processing, analysis and quantification

All data were recorded in a 5 x 5 or 6 x 6 field of view grid format. Data were first denoised using ‘advanced denoising’ in NIS-Analysis (Nikon), with a denoising power set to 0 for all channels. Background was corrected using a rolling ball algorithm (NIS-Analysis; Nikon), with a ball radius set to 0.96 μm. Grid images were next split to individual fields of view, which were subsequently corrected for drift using ‘align’ in NIS-Analysis. Regions of interest were selected by hand, cropped and, if needed, rotated using Fiji. Kymograms were generated using ‘montage’ in Fiji.

For intensity analysis during licensing in HSS, the intensity plots were generated in Fiji for individual molecules between 3 and 14 min of incubation time. Data were normalized to background (‘0’) and maximum intensity value (‘1’). Average intensity profiles were generated for each tested nucleosomal template, with a mean fluorescence value and standard deviation calculated at each time point. The mean value traces were then fitted to a single exponential decay model using Prism (GraphPad).

Replication fork velocities were calculated by measuring the distance travelled by an individual fork over time, in μm/min. Velocities were next converted to nt/min based on the measured average length of *λ* DNA from SYTOX staining. Mean fork velocities and associated standard deviations were calculated from a Gaussian fit to a histogram (GraphPad Prism).

For the analysis of fork-nucleosome collision outcomes, a number of criteria were implemented to ensure their reliable assignment and quantification. Only well-separated stretched *λ* molecules were included in the analysis. Histones that displayed thermal fluctuations inconsistent with the stretched *λ* DNA molecule were excluded from the analysis; for example, if a broken singly-tethered *λ* DNA is located close to a doubly-tethered *λ* DNA, its nucleosomes may at a first glance appear as part of the doubly-tethered molecule but are usually distinguishable through local fluctuations over time. Histone eviction was defined by the loss of histone fluorescence in the next time frame upon fork encounter. Histone transfer was assigned when, upon fork encounter, histone-associated fluorescence was incorporated into the replication bubble and could be followed for at least three subsequent time frames (3 minutes). Nucleosome (histone) sliding was determined by a unified histone-fork movement over at least 3 pixels (0.48 μM; ∼2.3 kbp). Replication fork stalling was assigned if a fork movement was arrested by a static (within 1 pixel) histone fluorescence for at least three time frames (3 minutes). Stalling events on nucleosomes showing particularly high histone fluorescence (over three times higher than the local average) were excluded from the analysis as they are likely to represent multiple nucleosomes on singly-tethered DNA or higher order local structure on doubly tethered DNA molecules. For the overall outcome quantification, all assigned events were counted, including the secondary events; for example, if a nucleosome (histone) sliding was followed by eviction, both sliding and eviction would be included in the quantification. A separate secondary outcome quantification was also conducted to gain insight into the outcome probability of nucleosome (histone) sliding and replication fork stalling.

## REFERENCES

Alabert, C., Barth, T.K., Reveron-Gomez, N., Sidoli, S., Schmidt, A., Jensen, O.N., Imhof, A., and Groth, A. (2015). Two distinct modes for propagation of histone PTMs across the cell cycle. Genes Dev 29, 585–590.

Alabert, C., Jasencakova, Z., and Groth, A. (2017). Chromatin Replication and Histone Dynamics. Adv Exp Med Biol 1042, 311–333.

Annunziato, A.T. (2013). Assembling chromatin: the long and winding road. Biochim Biophys Acta 1819, 196–210.

Annunziato, A.T. (2015). The Fork in the Road: Histone Partitioning During DNA Replication. Genes (Basel) 6, 353–371.

Bonev, B., and Cavalli, G. (2016). Organization and function of the 3D genome. Nat Rev Genet 17, 772.

Bowman, G.D. (2010). Mechanisms of ATP-dependent nucleosome sliding. Curr Opin Struct Biol 20, 73–81.

Clement, C., Orsi, G.A., Gatto, A., Boyarchuk, E., Forest, A., Hajj, B., Mine-Hattab, J., Garnier, M., Gurard-Levin, Z.A., Quivy, J.P., et al. (2018). High-resolution visualization of H3 variants during replication reveals their controlled recycling. Nat Commun 9, 3181.

Dyer, P.N., Edayathumangalam, R.S., White, C.L., Bao, Y., Chakravarthy, S., Muthurajan, U.M., and Luger, K. (2004). Reconstitution of nucleosome core particles from recombinant histones and DNA. Methods Enzymol 375, 23–44.

Escobar, T., Oksuz, O., Descostes, N., Bonasio, R., and Reinberg, D. (2019). Active and repressed chromatin domains exhibit distinct nucleosome segregation during DNA replication. bioRxiv, 418707.

Falk, M., Feodorova, Y., Naumova, N., Imakaev, M., Lajoie, B.R., Leonhardt, H., Joffe, B., Dekker, J., Fudenberg, G., Solovei, I., et al. (2019). Publisher Correction: Heterochromatin drives compartmentalization of inverted and conventional nuclei. Nature 572, E22.

Foltman, M., Evrin, C., De Piccoli, G., Jones, R.C., Edmondson, R.D., Katou, Y., Nakato, R., Shirahige, K., and Labib, K. (2013). Eukaryotic replisome components cooperate to process histones during chromosome replication. Cell Rep 3, 892–904.

Gan, H., Serra-Cardona, A., Hua, X., Zhou, H., Labib, K., Yu, C., and Zhang, Z. (2018). The Mcm2-Ctf4-Polalpha Axis Facilitates Parental Histone H3-H4 Transfer to Lagging Strands. Mol Cell 72, 140–151 e143.

Gasser, R., Koller, T., and Sogo, J.M. (1996). The stability of nucleosomes at the replication fork. J Mol Biol 258, 224–239.

Groth, A., Corpet, A., Cook, A.J., Roche, D., Bartek, J., Lukas, J., and Almouzni, G. (2007). Regulation of replication fork progression through histone supply and demand. Science 318, 1928–1931.

Gunjan, A., and Verreault, A. (2003). A Rad53 kinase-dependent surveillance mechanism that regulates histone protein levels in S. cerevisiae. Cell 115, 537–549.

Gupta, P., Zlatanova, J., and Tomschik, M. (2009). Nucleosome assembly depends on the torsion in the DNA molecule: a magnetic tweezers study. Biophys J 97, 3150–3157.

Gurard-Levin, Z.A., Quivy, J.P., and Almouzni, G. (2014). Histone chaperones: assisting histone traffic and nucleosome dynamics. Annu Rev Biochem 83, 487–517.

Han, M., Chang, M., Kim, U.J., and Grunstein, M. (1987). Histone H2B repression causes cell-cycle-specific arrest in yeast: effects on chromosomal segregation, replication, and transcription. Cell 48, 589–597.

Huang, H., Stromme, C.B., Saredi, G., Hodl, M., Strandsby, A., Gonzalez-Aguilera, C., Chen, S., Groth, A., and Patel, D.J. (2015). A unique binding mode enables MCM2 to chaperone histones H3-H4 at replication forks. Nat Struct Mol Biol 22, 618–626.

Ishimi, Y., Komamura, Y., You, Z., and Kimura, H. (1998). Biochemical function of mouse minichromosome maintenance 2 protein. J Biol Chem 273, 8369–8375.

Jackson, V. (1990). In vivo studies on the dynamics of histone-DNA interaction: evidence for nucleosome dissolution during replication and transcription and a low level of dissolution independent of both. Biochemistry 29, 719–731.

Kim, U.J., Han, M., Kayne, P., and Grunstein, M. (1988). Effects of histone H4 depletion on the cell cycle and transcription of Saccharomyces cerevisiae. EMBO J 7, 2211–2219.

Kimura, H., and Cook, P.R. (2001). Kinetics of core histones in living human cells: little exchange of H3 and H4 and some rapid exchange of H2B. J Cell Biol 153, 1341–1353.

Kose, H.B., Larsen, N.B., Duxin, J.P., and Yardimci, H. (2019). Dynamics of the Eukaryotic Replicative Helicase at Lagging-Strand Protein Barriers Support the Steric Exclusion Model. Cell Rep 26, 2113–2125 e2116.

Kurat, C.F., Yeeles, J.T.P., Patel, H., Early, A., and Diffley, J.F.X. (2017). Chromatin Controls DNA Replication Origin Selection, Lagging-Strand Synthesis, and Replication Fork Rates. Mol Cell 65, 117–130.

Lai, W.K.M., and Pugh, B.F. (2017). Understanding nucleosome dynamics and their links to gene expression and DNA replication. Nat Rev Mol Cell Biol 18, 548–562.

Larson, A.G., Elnatan, D., Keenen, M.M., Trnka, M.J., Johnston, J.B., Burlingame, A.L., Agard, D.A., Redding, S., and Narlikar, G.J. (2017). Liquid droplet formation by HP1alpha suggests a role for phase separation in heterochromatin. Nature 547, 236–240.

Lebofsky, R., Takahashi, T., and Walter, J.C. (2009). DNA replication in nucleus-free *Xenopus* egg extracts. Methods Mol Biol 521, 229–252.

Louters, L., and Chalkley, R. (1985). Exchange of histones H1, H2A, and H2B in vivo. Biochemistry 24, 3080–3085.

Loveland, A.B., Habuchi, S., Walter, J.C., and van Oijen, A.M. (2012). A general approach to break the concentration barrier in single-molecule imaging. Nat Methods 9, 987–992.

Luger, K., Mader, A.W., Richmond, R.K., Sargent, D.F., and Richmond, T.J. (1997). Crystal structure of the nucleosome core particle at 2.8 A resolution. Nature 389, 251–260.

MacAlpine, D.M., and Almouzni, G. (2013). Chromatin and DNA replication. Cold Spring Harb Perspect Biol 5, a010207.

Madamba, E.V., Berthet, E.B., and Francis, N.J. (2017). Inheritance of Histones H3 and H4 during DNA Replication In Vitro. Cell Rep 21, 1361–1374.

Marzluff, W.F., Wagner, E.J., and Duronio, R.J. (2008). Metabolism and regulation of canonical histone mRNAs: life without a poly(A) tail. Nat Rev Genet 9, 843–854.

Mejlvang, J., Feng, Y., Alabert, C., Neelsen, K.J., Jasencakova, Z., Zhao, X., Lees, M., Sandelin, A., Pasero, P., Lopes, M., et al. (2014). New histone supply regulates replication fork speed and PCNA unloading. J Cell Biol 204, 29–43.

Nelson, D.M., Ye, X., Hall, C., Santos, H., Ma, T., Kao, G.D., Yen, T.J., Harper, J.W., and Adams, P.D. (2002). Coupling of DNA synthesis and histone synthesis in S phase independent of cyclin/cdk2 activity. Mol Cell Biol 22, 7459–7472.

Newport, J., and Kirschner, M. (1982). A major developmental transition in early *Xenopus* embryos: II. Control of the onset of transcription. Cell 30, 687–696.

Petryk, N., Dalby, M., Wenger, A., Stromme, C.B., Strandsby, A., Andersson, R., and Groth, A. (2018). MCM2 promotes symmetric inheritance of modified histones during DNA replication. Science 361, 1389–1392.

Ramachandran, S., and Henikoff, S. (2015). Replicating Nucleosomes. Sci Adv 1.

Reinberg, D., and Vales, L.D. (2018). Chromatin domains rich in inheritance. Science 361, 33–34.

Reveron-Gomez, N., Gonzalez-Aguilera, C., Stewart-Morgan, K.R., Petryk, N., Flury, V., Graziano, S., Johansen, J.V., Jakobsen, J.S., Alabert, C., and Groth, A. (2018). Accurate Recycling of Parental Histones Reproduces the Histone Modification Landscape during DNA Replication. Mol Cell 72, 239–249 e235.

Rhind, N., and Gilbert, D.M. (2013). DNA replication timing. Cold Spring Harb Perspect Biol 5, a010132.

Sekinger, E.A., Moqtaderi, Z., and Struhl, K. (2005). Intrinsic histone-DNA interactions and low nucleosome density are important for preferential accessibility of promoter regions in yeast. Mol Cell 18, 735–748.

Serra-Cardona, A., and Zhang, Z. (2018). Replication-Coupled Nucleosome Assembly in the Passage of Epigenetic Information and Cell Identity. Trends Biochem Sci 43, 136–148.

Shundrovsky, A., Smith, C.L., Lis, J.T., Peterson, C.L., and Wang, M.D. (2006). Probing SWI/SNF remodeling of the nucleosome by unzipping single DNA molecules. Nat Struct Mol Biol 13, 549–554.

Sogo, J.M., Stahl, H., Koller, T., and Knippers, R. (1986). Structure of replicating simian virus 40 minichromosomes. The replication fork, core histone segregation and terminal structures. J Mol Biol 189, 189–204.

Solovei, I., Thanisch, K., and Feodorova, Y. (2016). How to rule the nucleus: divide et impera. Curr Opin Cell Biol 40, 47–59.

Stillman, B. (2018). Histone Modifications: Insights into Their Influence on Gene Expression. Cell 175, 6–9.

Strom, A.R., Emelyanov, A.V., Mir, M., Fyodorov, D.V., Darzacq, X., and Karpen, G.H. (2017). Phase separation drives heterochromatin domain formation. Nature 547, 241–245.

Swoboda, M., Henig, J., Cheng, H.M., Brugger, D., Haltrich, D., Plumere, N., and Schlierf, M. (2012). Enzymatic oxygen scavenging for photostability without pH drop in single-molecule experiments. ACS Nano 6, 6364–6369.

Thastrom, A., Bingham, L.M., and Widom, J. (2004a). Nucleosomal locations of dominant DNA sequence motifs for histone-DNA interactions and nucleosome positioning. J Mol Biol 338, 695–709.

Thastrom, A., Lowary, P.T., and Widom, J. (2004b). Measurement of histone-DNA interaction free energy in nucleosomes. Methods 33, 33–44.

Thiriet, C., and Hayes, J.J. (2006). Histone dynamics during transcription: exchange of H2A/H2B dimers and H3/H4 tetramers during pol II elongation. Results Probl Cell Differ 41, 77–90.

van Steensel, B., and Belmont, A.S. (2017). Lamina-Associated Domains: Links with Chromosome Architecture, Heterochromatin, and Gene Repression. Cell 169, 780–791.

Verreault, A., Kaufman, P.D., Kobayashi, R., and Stillman, B. (1998). Nucleosomal DNA regulates the core-histone-binding subunit of the human Hat1 acetyltransferase. Curr Biol 8, 96–108.

Walter, J., and Newport, J. (2000). Initiation of eukaryotic DNA replication: origin unwinding and sequential chromatin association of Cdc45, RPA, and DNA polymerase alpha. Mol Cell 5, 617–627.

Widom, J. (1998). Structure, dynamics, and function of chromatin in vitro. Annu Rev Biophys Biomol Struct 27, 285–327.

Woodland, H.R., Flynn, J.M., and Wyllie, A.J. (1979). Utilization of stored mRNA in *Xenopus* embryos and its replacement by newly synthesized transcripts: histone H1 synthesis using interspecies hybrids. Cell 18, 165–171.

Xu, M., Long, C., Chen, X., Huang, C., Chen, S., and Zhu, B. (2010). Partitioning of histone H3-H4 tetramers during DNA replication-dependent chromatin assembly. Science 328, 94–98.

Yardimci, H., Loveland, A.B., Habuchi, S., van Oijen, A.M., and Walter, J.C. (2010). Uncoupling of sister replisomes during eukaryotic DNA replication. Mol Cell 40, 834–840.

Yardimci, H., Loveland, A.B., van Oijen, A.M., and Walter, J.C. (2012). Single-molecule analysis of DNA replication in *Xenopus* egg extracts. Methods 57, 179–186.

Yu, C., Gan, H., Serra-Cardona, A., Zhang, L., Gan, S., Sharma, S., Johansson, E., Chabes, A., Xu, R.M., and Zhang, Z. (2018). A mechanism for preventing asymmetric histone segregation onto replicating DNA strands. Science 361, 1386–1389.

Zhou, K., Gaullier, G., and Luger, K. (2019). Nucleosome structure and dynamics are coming of age. Nat Struct Mol Biol 26, 3–13.

Zierhut, C., Jenness, C., Kimura, H., and Funabiki, H. (2014). Nucleosomal regulation of chromatin composition and nuclear assembly revealed by histone depletion. Nat Struct Mol Biol 21, 617–625.

