## Supplemental Information for "Single-molecule imaging reveals control of parental histone recycling by free histones during DNA replication"

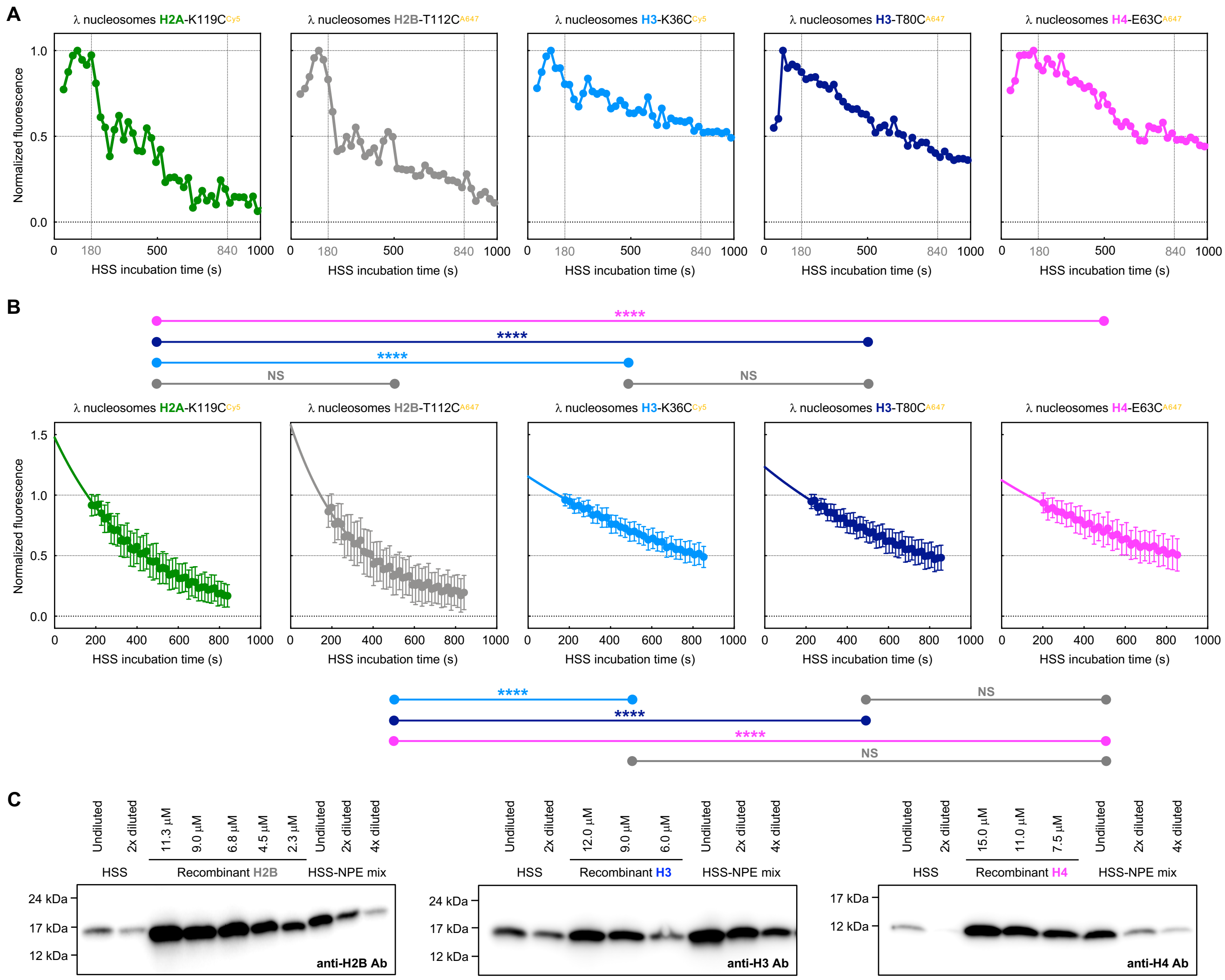


**Figure S1. Related to Figure 3. Histone dynamics during DNA licensing in HSS.**

(A) Representative intensity profiles for fluorescent λ nucleosomes during incubation in HSS. The profiles illustrate the initial increase in histone fluorescence, when HSS reaches the immobilized molecules (at 60-90sec) in the microfluidic device. The vertical dotted lines mark the range of data used for further intensity analysis presented in Figures 3D and S1B.

(B) Plots of the mean loss of fluorescent signal for λ nucleosomes during incubation in HSS. Data were fitted to a one phase decay model and the associated fitting parameters are listed in Table S2. P values extracted from the unpaired t test comparisons between data sets are presented above and below the plots.

(C) Western blots used to estimate the concentration of histone H2B, H3 and H4 in *Xenopus* egg extracts.

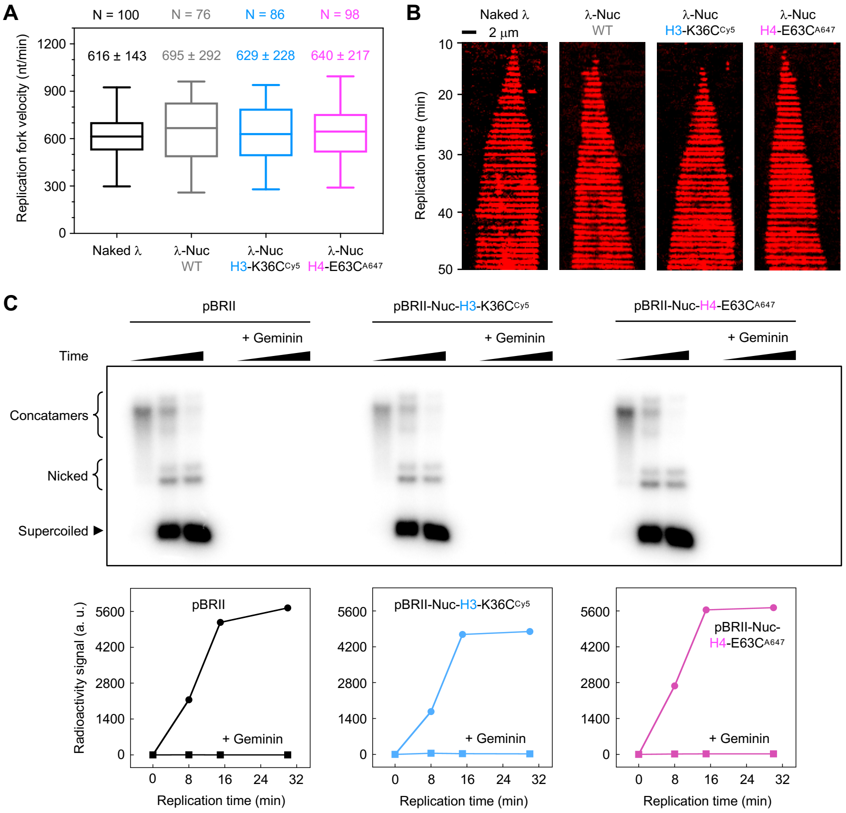


**Figure S2. Related to Figures 4, 5 and 6. Replication of λ nucleosomal templates in *Xenopus* egg extracts.**

(A) Box-and-whisker Tukey plot of replication fork velocities for naked λ and λ nucleosomes containing wild-type octamer, octamer labelled at H3-K36C$^{Cy5}$ and octamer labelled at H4-E63C$^{A647}$. Velocities were calculated from real-time single-molecule experiments. Values above the box plots indicate the mean replication fork velocity extracted from the Gaussian fit, plus and minus standard deviation. The number of values analyzed per data set (N) is also shown.

(B) Kymograms of Fen1-KikGR fluorescence indicating growth of replication bubble over time.

(C) Bulk replication assay for naked pBRII plasmid and pBRII plasmid containing nucleosomes labelled at H3-K36C$^{Cy5}$ or H4-E63C$^{A647}$. For each replicated template, a negative control is also presented (+ Geminin). Quantification of the replication efficiency, as measured by the radioactivity signal, is presented in the lower panel.

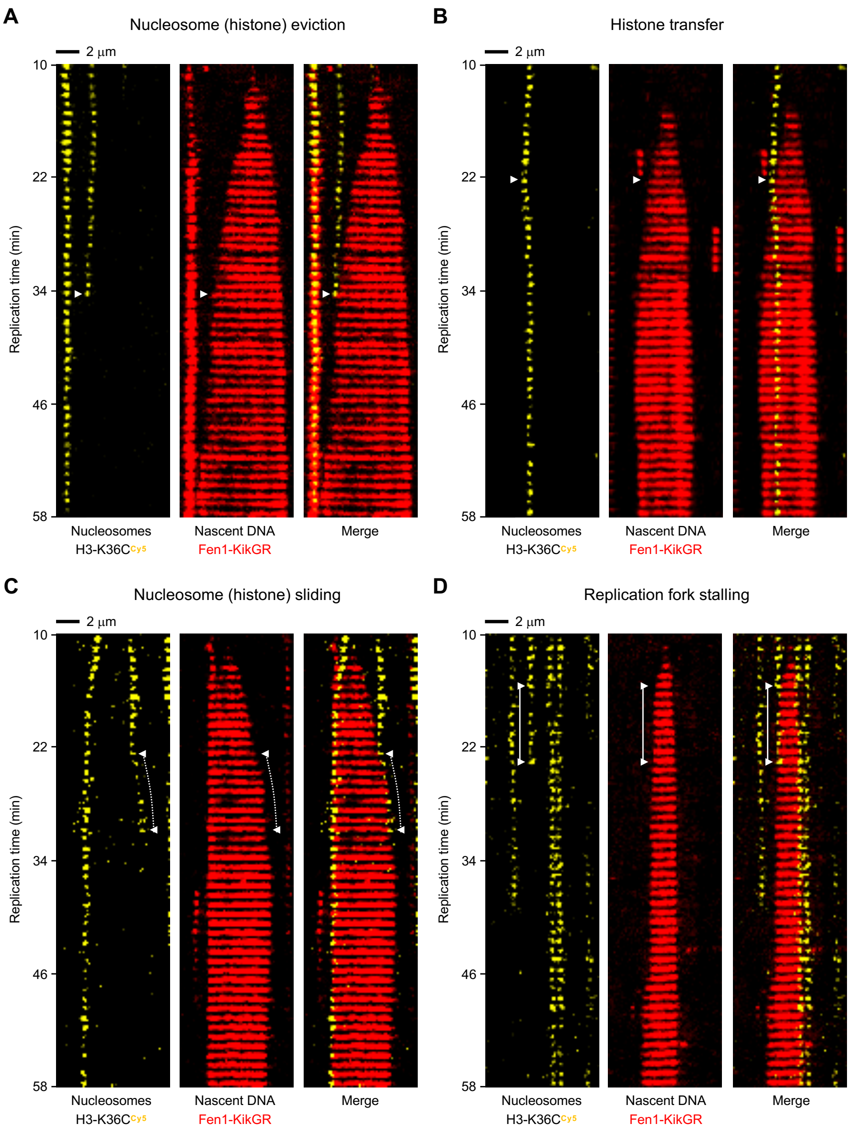


**Figure S3. Related to Figures 4, 5 and 6. Outcomes of the replication fork collision with nucleosomes containing H3-K36C$^{Cy5}$ during DNA replication in *Xenopus* egg extracts.** For each specified outcome, data are presented as kymograms of nucleosome-associated fluorescence (H3-K36C$^{Cy5}$; yellow; left panels), Fen1-KikGR signal indicating nascent DNA (red; central panels) and both signals together (merge; right panels). Time and size scales are presented. The white triangles mark the point of initial encounter between the replication fork and nucleosome. Dotted lines indicate sliding events, whereas solid lines correspond to replication fork stalling.

(A) Nucleosome (histone) eviction.

(B) Histone transfer onto daughter strands.

(C) Nucleosome (histone) sliding terminating in nucleosome (histone) eviction.

(D) Replication fork stalling terminating in nucleosome (histone) eviction.

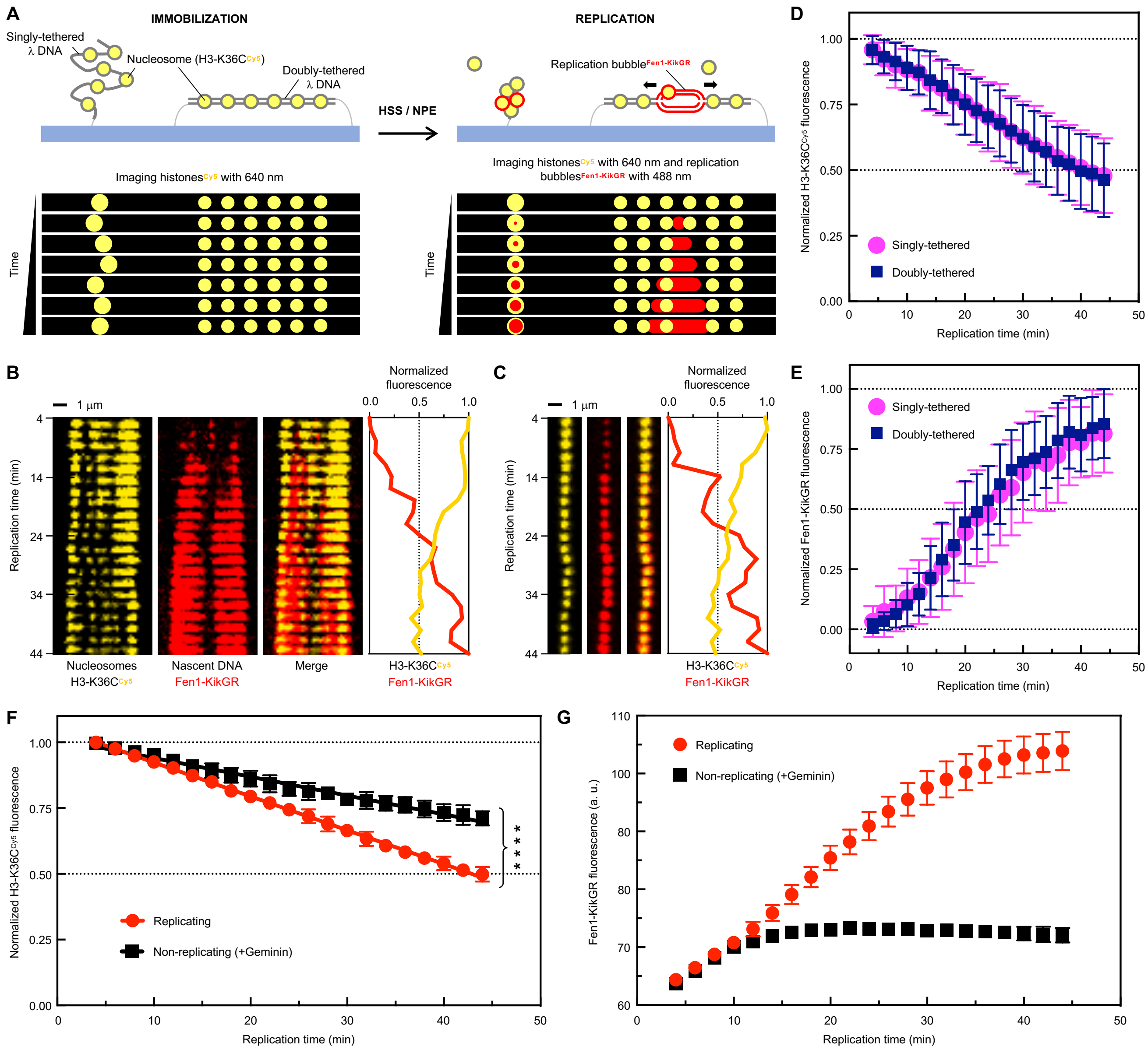


**Figure S4. Related to Figure 6. Replication of singly- and doubly-tethered λ nucleosomes labelled at H3-K36C$^{Cy5}$ in *Xenopus* egg extracts.**

(A) Schematic of the experimental set-up. Doubly- and singly-biotinylated λ DNA molecules containing fluorescent nucleosomes are attached to the surface so that approximately 50% of the molecules are tethered in a stretched form while the remaining molecules are tethered at only one end. Doubly-tethered λ nucleosomes display the characteristic 'beads-on-a-string' appearance, whereas singly-tethered λ nucleosomes show up as a spot of fluorescence that can be stretched under buffer flow to reveal the individual 'beads' (Video 1). The immobilized DNA is licensed in high-speed supernatant (HSS), which causes the singly-tethered molecules to compact (they no longer stretch under flow) due to the deposition of extract proteins. Replication is initiated upon introduction of nucleoplasmic extract (NPE) supplemented with a fluorescent fusion protein Fen1-KikGR, which decorates replication bubbles. Firing was unrestricted in these experiments. Cy5-labelled histones and Fen1-KikGR are imaged with 640-nm and 488-nm laser, respectively. To minimize photobleaching, images were taken at two-minute intervals. Replication of singly-tethered λ nucleosomes is manifested by the appearance and subsequent increase of Fen1-KikGR fluorescence, colocalized with the H3-K36C$^{Cy5}$ signal.

(B and C) Kymograms and corresponding intensity profiles for doubly- (B) and singly-tethered (C) λ nucleosomes. Kymograms of H3-K36C$^{Cy5}$ fluorescence (yellow; left panels), Fen1-KikGR (red; central panels) and both signals together (merge; right panels) are presented. Time and size scales are indicated.

(D) Plot showing the mean loss of H3-K36C$^{Cy5}$ fluorescence for doubly- (blue squares) and singly-tethered (magenta circles) λ nucleosomes during replication under unrestricted firing conditions. 110 molecules were analyzed for each data set. Individual fluorescence decay traces were normalized to background ('0') and maximum values of fluorescence ('1'). A mean fluorescence value and standard deviation were calculated and plotted for each time point. We observed no difference in the loss of H3-K36C$^{Cy5}$ fluorescence between the doubly- and singly-tethered λ nucleosomes.

(E) Plot showing the mean increase of Fen1-KikGR fluorescence for doubly- (blue squares) and singly-tethered (magenta circles) λ nucleosomes during replication under unrestricted firing conditions. 110 molecules were analyzed for each data set. Individual fluorescence decay traces were normalized to minimum ('0') and maximum values of fluorescence ('1'). A mean fluorescence value and standard deviation were calculated and plotted for each time point. We observed no difference in the firing timing between the doubly- and singly-tethered λ nucleosomes.

(F) Plot showing the mean loss of H3-K36C$^{Cy5}$ fluorescence for 1:1 mixture of doubly- and singly-tethered λ nucleosomes during replication under unrestricted firing conditions (red circles) versus non-replicating control (black squares; +Geminin). 36 fields of view were analyzed for each data set. Decay traces for individual fields of view were normalized to background ('0') and maximum values of fluorescence ('1'). A mean fluorescence value and standard deviation were calculated and plotted for each time point. Data sets were fitted to a linear regression model (see Table S3 for fitting parameters) and the statistical significance of the differences between the slopes was estimated as <0.0001. The loss of histone-associated fluorescence is significantly faster for replicating nucleosomes than for nucleosomes incubated in non-replicating extracts (see panel G for the replication efficiency comparison). Based on this analysis we conclude that the observed losses of histones during replication are largely caused by active replication forks, with some contribution from replication-independent histone exchange and photobleaching.

(G) Plot showing the mean increase of Fen1-KikGR fluorescence for 1:1 mixture of doubly- and singly-tethered λ nucleosomes during replication under unrestricted firing conditions (red circles) versus non-replicating control (black squares; +Geminin). 36 fields of view were analyzed for each data set. A mean fluorescence value and standard deviation were calculated and plotted for each time point.

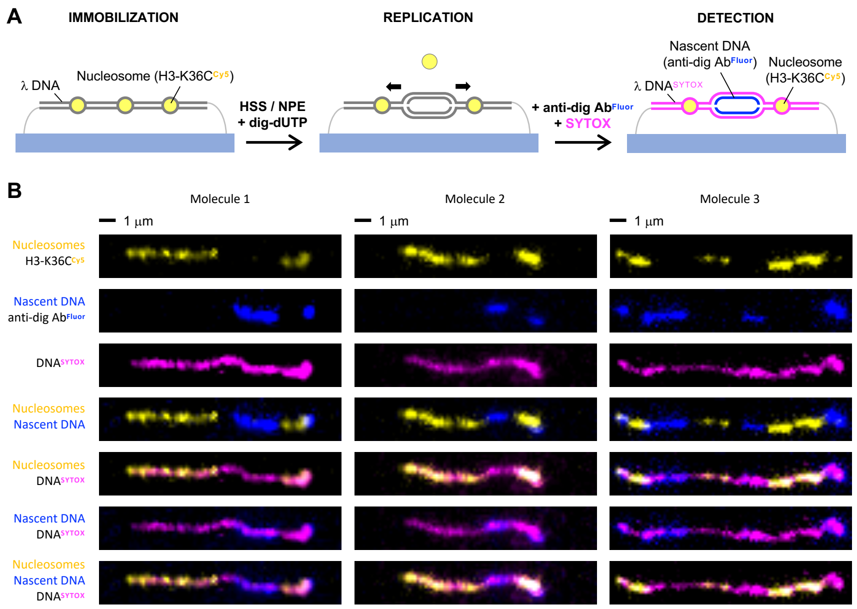


**Figure S5. Related to Figure 6. Replication of doubly-tethered λ nucleosomes labelled at H3-K36C$^{Cy5}$ in *Xenopus* egg extracts in the absence of Fen1-KikGR.**

(A) Schematic of the experimental set-up. Doubly-biotinylated λ DNA molecules containing fluorescent nucleosomes are stretched under flow and attached to the surface. The immobilized DNA is licensed in high-speed supernatant (HSS) for 15 minutes. Replication is initiated upon introduction of nucleoplasmic extract (NPE) supplemented with digoxygenin-11-dUTP (dig-dUTP). After 30 minutes, replication is stopped by flowing in buffer containing 20 mM Tris pH 7.5, 10 mM EDTA and 0.5 M NaCl. Under these conditions, extracts are washed out from the flow cell but nucleosomes remain intact on the immobilized DNA and can be imaged with 640-nm laser. Nascent DNA, containing dig-dUTP incorporated during replication, is visualized through immunostaining with an anti-digoxygenin antibody labelled with fluorescein (anti-dig Ab$^{Fluor}$; excited with 488-nm laser). In addition, non- and replicated DNA is stained with SYTOX Orange and visualized using 561-nm laser.

(B) Examples of λ molecules containing H3-K36C$^{Cy5}$ nucleosomes replicated in the absence of Fen1-KikGR. Post-replication detection of H3-K36C$^{Cy5}$ nucleosomes (yellow), nascent DNA (blue; anti-dig Ab$^{Fluor}$) and overall DNA (magenta; SYTOX Orange) demonstrates that nucleosome-free zones colocalize with nascent DNA tracts, indicating that H3-K36C$^{Cy5}$ histones do not efficiently transfer behind the replication fork in the absence of Fen1-KikGR. Note that the high salt wash, required for replication termination in this assay, leads to non-specific sticking of the immobilized DNA molecules to the surface, which appear bent rather than straight lines under the microscope.

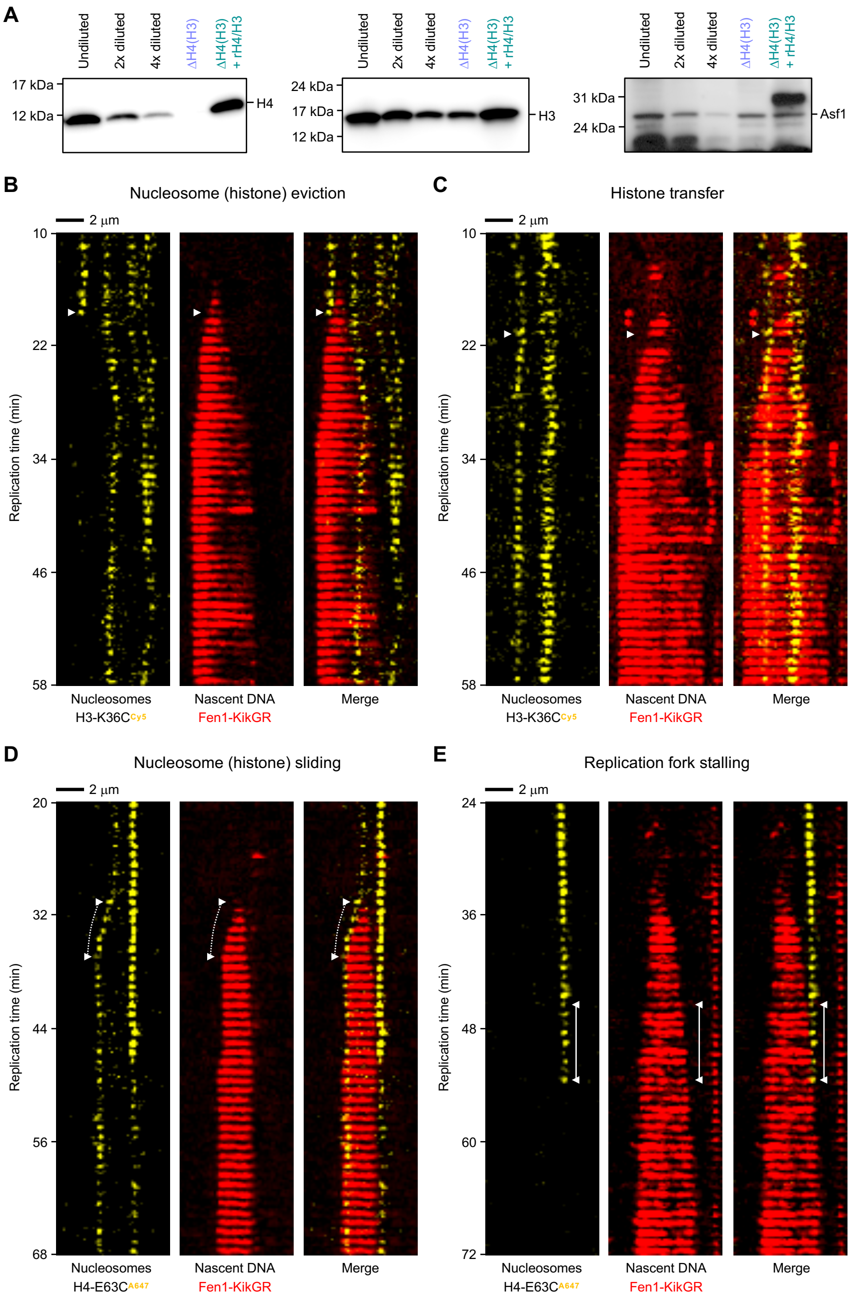


**Figure S6. Related to Figure 6. Outcomes of the replication fork collision with nucleosomes during DNA replication in *Xenopus* egg extracts depleted of histones H3 and H4.** For each specified outcome, data are presented as kymograms of nucleosome-associated fluorescence (H3-K36C$^{Cy5}$ or H4-E63C$^{A647}$; yellow; left panels), Fen1-KikGR signal indicating nascent DNA (red; central panels) and both signals together (merge; right panels). Time and size scales are presented. The white triangles mark the point of initial encounter between the replication fork and nucleosome. Dotted lines indicate sliding events, whereas solid lines correspond to replication fork stalling.

(A) Western blots used to estimate the levels of histones H4 (left panel) and H3 (central panel), as well as the Asf1 histone chaperone (right panel) in regular *Xenopus* egg extracts (undiluted, 2x diluted, 4x diluted), extracts depleted of histone H4 and H3 (ΔH4/H3) and depleted extracts supplemented with recombinant histones H3 and H4 (ΔH4/H3 + rH4/H3).

(B) Nucleosome (histone) eviction.

(C) Histone transfer onto daughter strands.

(D) Nucleosome (histone) sliding is followed by replication fork stalling, which leads to histone transfer onto daughter strands.

(E) Replication fork stalling terminating in nucleosome (histone) eviction.

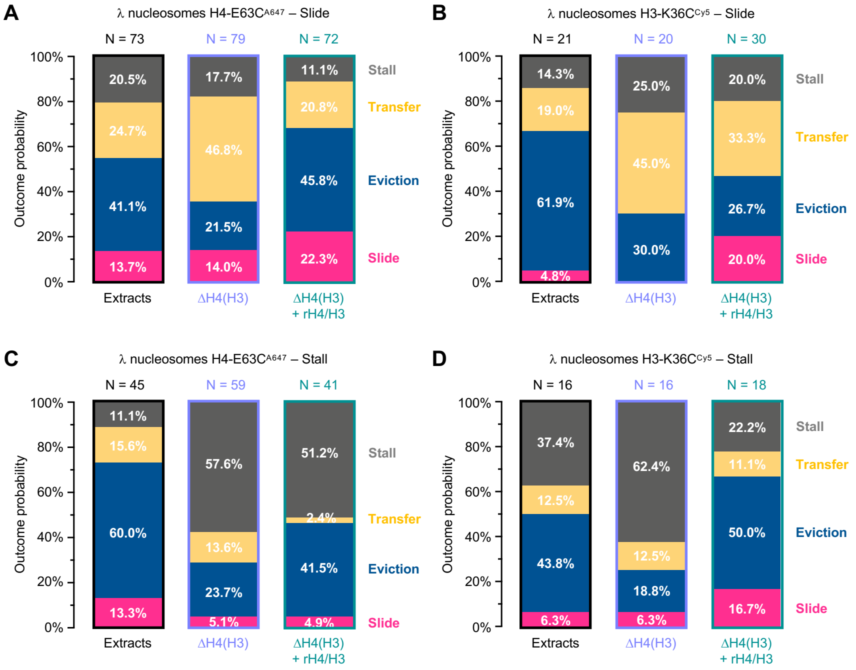


**Figure S7. Related to Figure 6. Quantification of the secondary outcomes of replication fork collision with nucleosomes.** Collisions were analyzed for assays conducted in regular undepleted extracts (black borders; left panels), extracts depleted of histones H4 and H3 (blue borders; central panels) and extracts depleted of endogenous histones but supplemented with recombinant H4 and H3 (green borders; right panels). N indicates the total number of analyzed collisions. For each condition, data from at least two biological repeats were pooled in the analysis.

(A) Quantification of the outcome of nucleosome/histone sliding for λ nucleosomes containing H4-E63C$^{A647}$.

(B) Quantification of the outcome of nucleosome/histone sliding for λ nucleosomes containing H3-K36C$^{Cy5}$.

(C) Quantification of the outcome of replication fork stalling for λ nucleosomes containing H4-E63C$^{A647}$.

(D) Quantification of the outcome of replication fork stalling for λ nucleosomes containing H3-K36C$^{Cy5}$.

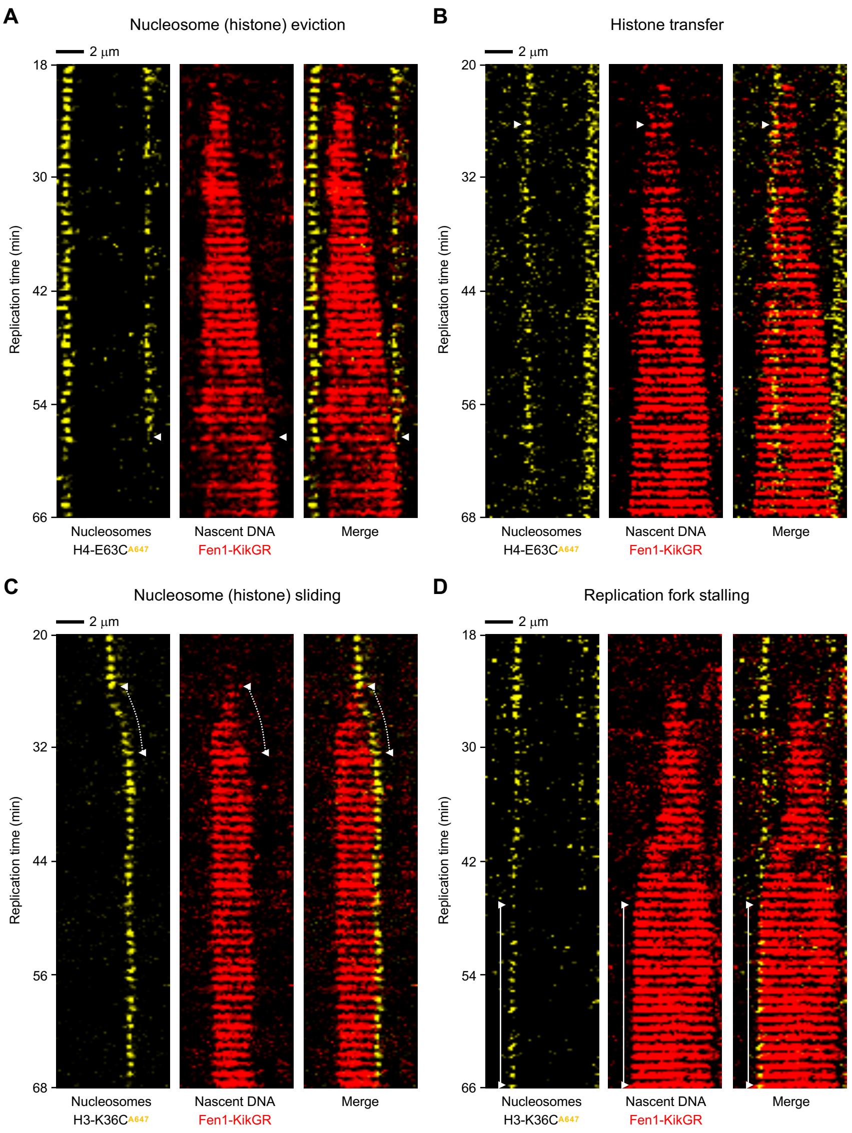


**Figure S8. Related to Figure 6. Outcomes of the replication fork collision with nucleosomes during DNA replication in *Xenopus* egg extracts depleted of endogenous histones H3 and H4, and supplemented with recombinant histones H3 and H4.** For each specified outcome, data are presented as kymograms of nucleosome-associated fluorescence (H3-K36C$^{Cy5}$ or H4-E63C$^{A647}$; yellow; left panels), Fen1-KikGR signal indicating nascent DNA (red; central panels) and both signals together (merge; right panels). Time and size scales are presented. The white triangles mark the point of initial encounter between the replication fork and nucleosome. Dotted lines indicate sliding events, whereas solid lines correspond to replication fork stalling.

(A) Nucleosome (histone) eviction.

(B) Histone transfer onto daughter strands.

(C) Nucleosome (histone) sliding followed by replication fork stalling.

(D) Replication fork stalling.

**Table S1. Related to Figure 1. Molecular weight of proteins used in this work**

| | Molecular weight (Da) | |
|---|---|---|
| **Protein** | **Theoretical** | **Mass spectrometry** |
| H2A | 13 950.2 | 13 949.6 |
| H2A-K119C | 13 925.1 | 13 924.6 |
| H2B | 13 493.7 | 13 493.1 |
| H2B-T112C | 13 495.7 | 13 495.0 |
| H3 | 15 270.8 | 15 270.2 |
| H3-K36C | 15 245.8 | 15 245.0 |
| H3-T80C-C110A | 15 240.8 | 15 240.2 |
| H4 | 11 236.1 | 11 235.7 |
| H4-E63C | 11 210.1 | 11 209.5 |

**Table S2. Related to Figures 3 and S1. Fitting parameters for histone dynamics data in HSS**

| **Protein** | **H2A-K119C$^{Cy5}$** | **H2B-T112C$^{A647}$** | **H3-K36C$^{Cy5}$** | **H3-T80C$^{A647}$** | **H4-E63C$^{A647}$** |
|---|---|---|---|---|---|
| N | 127 | 128 | 122 | 133 | 132 |
| $Y_0$ | $1.476 \pm 0.022$ | $1.580 \pm 0.039$ | $1.154 \pm 0.011$ | $1.232 \pm 0.015$ | $1.123 \pm 0.018$ |
| Plateau | $-0.023 \pm 0.018$ | $0.102 \pm 0.012$ | $-0.094 \pm 0.089$ | $0.012 \pm 0.060$ | $-0.008 \pm 0.129$ |
| Amplitude | $1.500 \pm 0.012$ | $1.478 \pm 0.030$ | $1.248 \pm 0.079$ | $1.220 \pm 0.047$ | $1.131 \pm 0.112$ |
| $K$ ($s^{-1}$) | $(2.417 \pm 0.091) \times 10^{-3}$ | $(3.448 \pm 0.138) \times 10^{-3}$ | $(0.894 \pm 0.099) \times 10^{-3}$ | $(1.148 \pm 0.104) \times 10^{-3}$ | $(0.935 \pm 0.172) \times 10^{-3}$ |
| $t_{0.5}$ (s) | 286.7 | 201.0 | 775.0 | 603.8 | 741.2 |
| $R^2$ | 0.7683 | 0.6161 | 0.7725 | 0.7168 | 0.5362 |

N indicates the number of individual decay traces used to generate plots of the mean loss of fluorescence.
Mean loss of fluorescence data were fitted to a one phase decay model, where $Y = (Y_0 - \text{Plateau})*\exp(-K*X) + \text{Plateau}$.
$Y_0$ is the Y value when X (time) is zero, $K$ is the rate constant and $t_{0.5}$ is the half-life, calculated as $\ln(2)/K$.

**Table S3. Related to Figure S4F. Fitting parameters for the loss of histone fluorescence data for replicating and non-replicating λ nucleosomes**

| Protein | Replicating | Non-replicating |
|---|---|---|
| Slope ($min^{-1}$) | -0.01288 ± 0.00007 | -0.007149 ± 0.00009 |
| Y-intercept | 1.052 ± 0.002 | 1.012 ± 0.002 |
| X-intercept (min) | 81.72 | 141.6 |
| 1/slope (min) | -77.66 | -139.9 |
| $R^2$ | 0.9787 | 0.9002 |

Data were fitted to a linear regression model, where Y = Slope* X + Y-intercept. In this case, the slope indicates the rate constant of the histone fluorescence loss.
